# Biomolecular condensates govern PARP inhibitor trapping and present mechanisms of resistance

**DOI:** 10.1101/2022.04.27.489674

**Authors:** Angelica A. Gopal, Bianca Fernandez, Justin Delano, Ralph Weissleder, J. Matthew Dubach

## Abstract

Poly(ADP-ribose) polymerase (PARP) inhibitors (PARPi) are a class of cancer drugs that enzymatically inhibit PARP activity at sites of DNA damage. In the context of BRCA mutations, PARPi can be synthetically lethal, presenting ideal genetic targeting. Yet, PARPi function primarily by trapping PARP1 onto sites of DNA damage. How PARPi trap and why some are better trappers remain unknown. Here, we show trapping occurs primarily through a kinetic phenomenon within biomolecular condensates that correlates with PARPi k_off_. Our results suggest PARP trapping is not the physical stalling of PARP1 on DNA, rather the high probability of PARP re-binding damaged DNA in the absence of other DNA binding protein recruitment. Furthermore, we found recruitment of the DNA binding protein RPA1 correlates to cell line PARPi sensitivity independent of RPA1 expression, demonstrating that condensate recruitment alone can impact efficacy. These results shed new light on how PARPi function, describe how PARPi properties correlate to trapping potency, and suggest previously unknown mechanisms of PARPi resistance.

---

PARP1 is an abundant nuclear protein^1^ that rapidly binds damaged DNA^2^ becoming active through allosteric conformational changes^3^. Active PARP1 produces prolific poly(ADP-ribose) (PAR) modification of thousands of proteins^4-6^ to recruit DNA damage response (DDR) proteins^7^, thus beginning the DNA damage response. All PARPi target the NAD^+^ binding pocket to enzymatically inhibit PAR production^8^ and several have reached the clinic^9-11^. But, loss of PARP1 does not impact cellular survival as much as PARPi treatment^12-14^, revealing that PARPi work largely by “trapping” PARP1 onto DNA. However, some PARPi are much more potent trappers than others and subsequently much more potent drugs^12,13^, despite having binding affinities within an order of magnitude.

PARP trapping, the primary driver of PARPi efficacy^15^, directly correlates to cellular response in HT1080 cells (Fig. 1a and Fig. S1a & b). Under the prevailing trapping theory - PARPi prevent PARP1 dissociation from DNA^16^ - it remains unclear why some PARPi are better trappers, and thus much more potent drugs through on-target effects (Fig. 1a and Fig. S1c-e). Elaborate measurements of PARP1 binding to DNA show altered affinities when PARP1 is engaged to a PARPi through reverse allosteric protein structure shifts^17^. However, the induced affinities of clinical PARPi do not correlate with observed trapping (Table S1, Fig. 1a). Additionally, PARP trapping also occurs through purely enzymatic mechanisms. Reduction in cellular NAD^+^ concentration (the PARP1 substrate) through nicotinamide phosphoribosyltransferase (NAMPT) inhibition alone induces trapping^18^ (Fig. 1b & Fig. S1f & g). But, PARP trapping is not canonically enzymatic. Veliparib at 100 μM is unable to reproduce the trapping observed with just 1 μM talazoparib, despite having a PARP1 affinity only one order of magnitude lower (Table S1 and Fig. S1b). Therefore, trapping (the PARPi-PARP1-DNA complex) is hypothesized to arise from a combination of PARPi reverse allosteric impact on PARP1 affinity for DNA^17^ and enzymatic inhibition, which limits PAR production, preventing self-PARylation induced release from DNA^12,13^ (Fig. 1d).

**Fig. 1.**
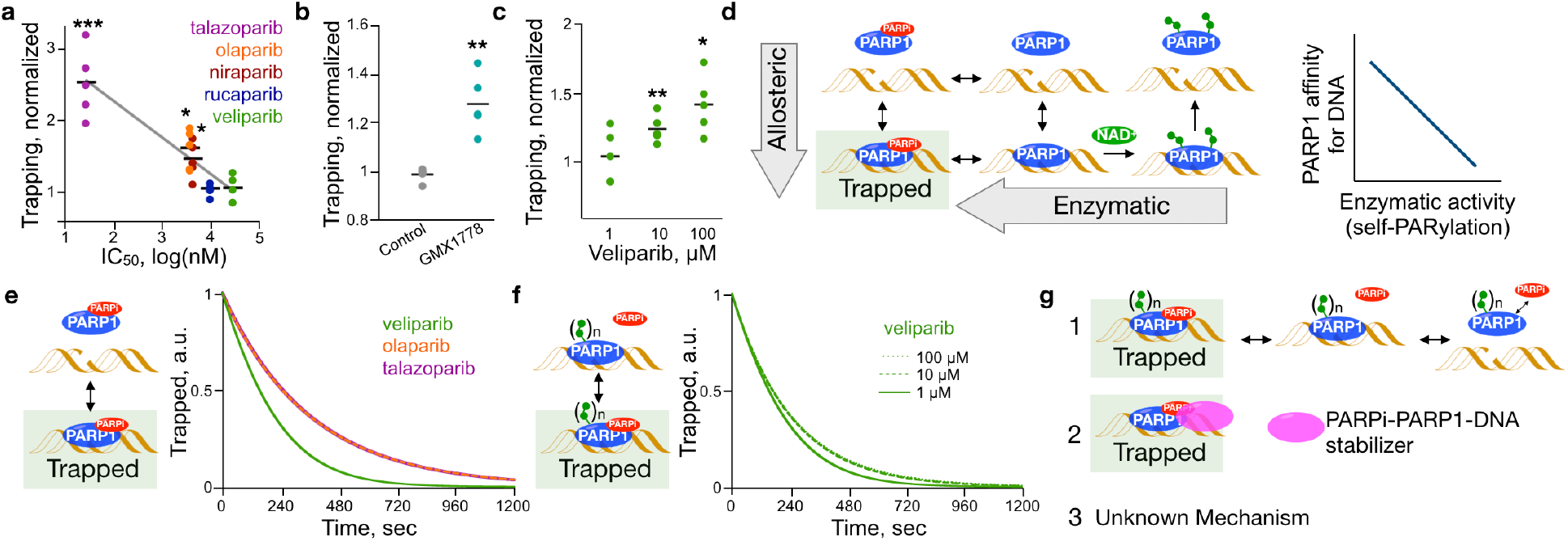
PARP trapping is not driven by the physical engagement of PARP1 to DNA. (**a**) Normalized PARP1 trapping in HT1080 cells treated with PARPi (1 μM overnight) versus IC_50_ with linear fit (gray line), individual measurements, n = 5 with average (black line). (**b**) Normalized trapping in HT1080 cells or cells treated with 10 μM GMX1778 overnight, n = 5 with average (black line). (**c**) Normalized trapping in HT1080 cells treated with veliparib overnight, n = 5 with average (black line). (**d**) Allosteric shifts in PARP1 affinity for DNA or PARPi enzymatic inhibition are thought to increase trapping and expected dependency of PARP1 affinity for DNA on self-PARylation. (**e-f**) ODE solution for the amount of trapped PARP1 e) in the presence of PARPi and omitting PAR-induced release, f) with different veliparib concentrations. (**g**) Potential mechanisms driving cellular PARP trapping. All data, * p < 0.005, ** p < 0.001, *** p < 0.0001 vs. control (Student’s t test).

We tested the trapping hypothesis through a set of ordinary differential equations (ODE) (Supplemental Data and Fig. S2 & 3) using established binding constants (Table S1). Considering allosteric impact alone, the duration of PARP1 bound to DNA is entirely dependent on the dissociation constant of PARP1-PARPi from DNA (Fig. 1e and Table S1). Therefore, based on previous measurements^17^, olaparib-bound PARP1 has the longest DNA interaction, which is both shorter than observed cellular trapping^17,19^ and does not agree with established differential PARPi trapping potency (Fig. 1a). We then calculated the enzymatic impact on PARP1-DNA residency through the presence of veliparib at increasing concentrations. Here we sought to test if increasing self-PARylation could impact self-PARylation induced release. However, the complex duration was driven largely by the dissociation of PARP1 from DNA with slight increases in duration at higher veliparib concentrations (Fig. 1f), which failed to capture the differences observed in cells (Fig. 1c).

Since the dissociation constant of PARP1 from DNA governs the canonical PARP trapping hypothesis we reasoned that PARP trapping must be driven by something else. We arrived at three different possible mechanisms (Fig. 1g). First, trapping must involve release and rebinding of PARP1 and self-PARylation accumulates based on enzymatic activity, reducing PARP1 affinity for DNA. Previous *in vitro* measurements show PARP1-DNA interactions lasting several hours in the presence of PARPi^12,13^, and correlate to differential PARPi trapping observed in cells. These previous findings indicate that PARP1 dissociates and rebinds DNA numerous times before sufficient self-PARylation is accumulated to prevent rebinding. However, to be correct, this mechanism relies on the same PARP1 molecule or molecules rebinding DNA and accumulating sufficient self-PARylation. Yet, the recent observation that PARP1 exchanges at sites of DNA damage independent of the presence of PARPi^19^ suggests that this is not possible, and it is unclear how unbound PARylated PARP1 would not diffuse away or be replaced by an unPARylated PARP1. Therefore, we removed this mechanism from consideration. A second mechanism could be the presence of cellular factors that drive increased PARP1-DNA interaction in a PARPi-specific manner. This mechanism would be similar to the presence of HPF1 interacting with PARP1 to drive preferential serine PARylation^20^. Lastly, there could be an unknown mechanism driving PARP trapping.

We first sought to determine if the PARPi-PARP1-DNA complex is altered in cells. We previously found that the equilibrium binding affinity inside cells for three PARPi was consistent with *in vitro* measurements^21^. Yet, we reasoned that if an extended duration of the PARPi-PARP1-DNA complex in the cellular environment drives trapping, the dissociation constant (k_off_) of PARPi from PARP1 would also be extended in a PARPi-specific manner. We developed an assay to quantify inhibitor k_off_ in live cells (Fig. 2a, Fig. S4 and Supplemental Data), measuring fluorescence anisotropy^21^ (Fig. 2b) of a fluorescently labeled PARPi, olaparib covalently attached to BODIPY FL^22^, in competitive binding experiments. We observed significantly different k_off_ rates for veliparib, olaparib and talazoparib in HT1080 cells (Fig. 2c and d), with values similar to those previously determined in purified protein (Table S1), indicating the duration of the PARPi-PARP1-DNA complex is not extended in cells. However, we found correlation between observed k_off_ rates and cell-type basal PARP activity in different cell lines (Fig. S5a & b), contrary to the equilibrium binding affinity^21^. This PARP activity impact was observed in a HT1080 cells alone by activating PARP through addition of the DNA alkylating agent temozolomide (TMZ), which causes broad DNA damage repair pathway activation^23^ (Fig. S5c). However, all three PARPi k_off_ values showed a PARP activation dependency (Fig. 2e), with a response expected from the integration of two distinct k_off_ values for each PARPi basal value (Fig. S5d-f). Thus, we concluded that the observed dissociation of PARPi from PARP was slower when PARP was activated, which is in contrast to previous measurements outside cells^18^, but this dependency was not unique to higher trapping PARPi. We also found the association binding constant (k_on_) of fluorescent olaparib was significantly lower when PARP was activated (Fig. S6a and c). We modeled the impact of k_on_ on k_off_ measurements and found a lower k_on_ does not fully explain the observed k_off_ dependency on PARP activation (Fig. S6b and d), yet the shifts in k_on_ and k_off_ are similar. These results indicate there is no PARPi-specific stabilization of the PARPi-PARP1-DNA complex in cells, but all binding rates are lower when PARP is activated.

**Fig. 2.**
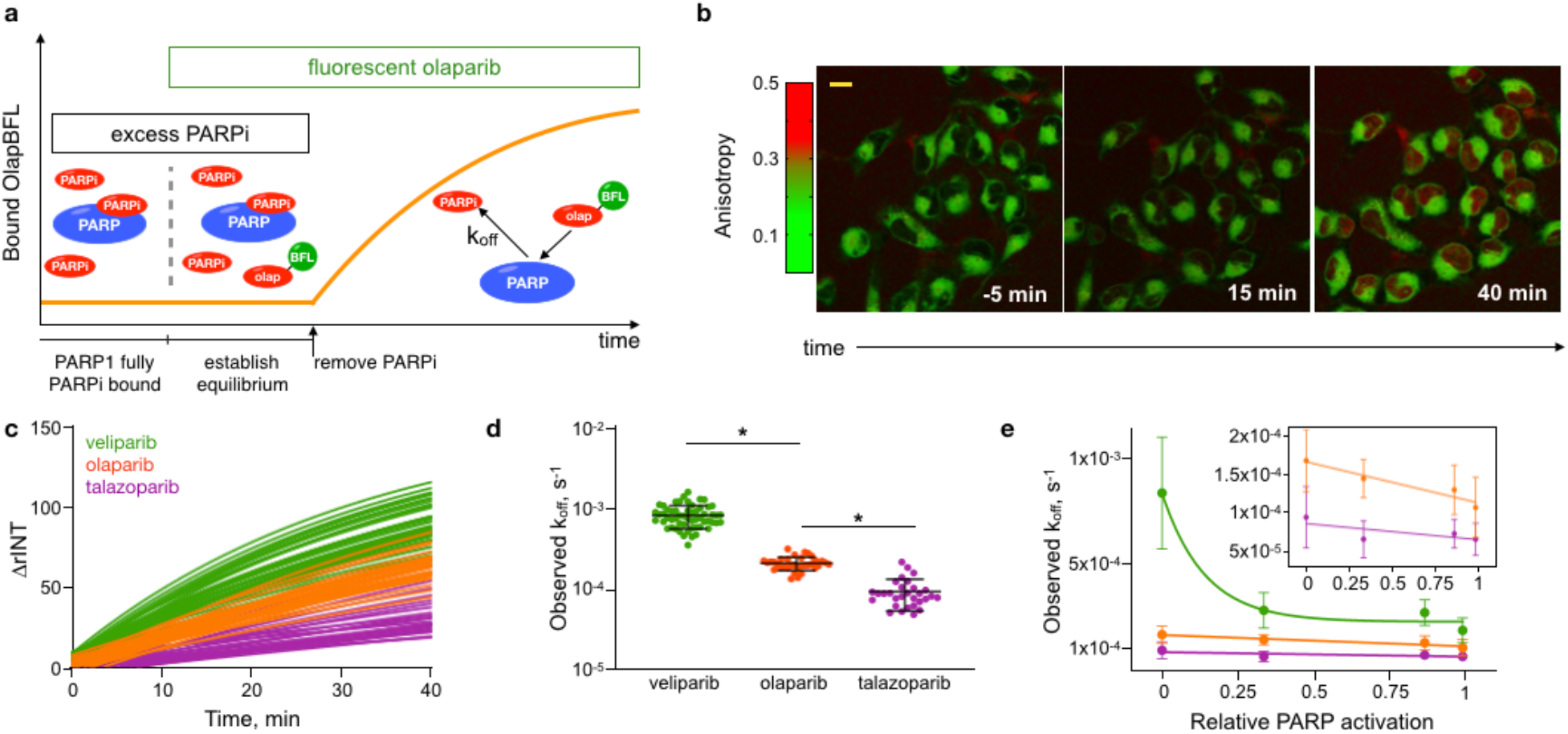
The PARPi-PARP1-DNA complex is not stabilized in cells but is PARP1 activity dependent. (**a**) An assay to measure dissociation of bound drug from cellular PARP. Excess PARPi occupies PARP target inside cells, fluorescently labeled olaparib (olap, BFL = BODIPY FL) is then added and PARPi is removed. The binding rate of fluorescent drug represents the dissociation (k_off_) of PARPi. (**b**) Representative anisotropy images of fluorescent olaparib in HT1080 cells during dissociation measurements, PARPi is removed at t = 0, scale bar = 5 μM. (**c**) ΔrINT measurements of fluorescent olaparib binding in the dissociation assay for 3 different PARPi, shown are fitted one phase association curves for single cells. (**d**) Single cell observed dissociation constants (k_off_) for 3 different PARPi, shown are individual cells with average and st. dev., n=30-57 cells, >= 4 biological repeats, * p < 1×10^−16^ (Student’s t test). (**e**) Observed k_off_ as a function of relative PARP activity for 3 PARPi, data are average with st. dev., n >= 6 cells, and fitted one phase dissociation curve.

One possible explanation for our observed PARP activation impact on all binding rates is an altered environment. Therefore, we turned our attention to a hallmark of PARP activity – the formation of DNA damage biomolecular condensates through PAR production and protein recruitment^24-29^. It is well established that PARPi alter protein recruitment^30-32^ and we hypothesized that altered recruitment would enable PARP dissociation from and rebinding to damaged DNA by preventing recruited proteins from displacing PARP. We reasoned that reduced PAR-based recruitment would produce a higher condensate density of PAR-independent markers with a correlation to PARP trapping (Fig. 3a & b). To quantify density within condensates we used segmented image correlation spectroscopy (ICS)^33^ of whole nuclei on immunofluorescence (IF) images of γH_2_AX, a marker of double-strand DNA breaks^34^. Here, ICS measures the average aggregation state of γH_2_AX within the nucleus. PARPi treatment overnight treatment at 1 μM induced condensate formation of γH_2_AX in HT1080 cells (Fig. 3c), with ICS degree of aggregation (DA) values correlating to trapping across 5 different PARPi (Fig. 3d). However, γH_2_AX levels, measured by intensity, did not have as strong a correlation to PARPi trapping (Fig. S7a). We also found γH_2_AX DA correlated to veliparib concentration dependent trapping (Fig. 3e). And, NAMPT inhibition with GMX1778 alone significantly increased γH_2_AX DA, correlating to measured trapping (Fig. 3f), despite no significant difference in γH_2_AX levels (Fig. S7c & d).

**Fig. 3.**
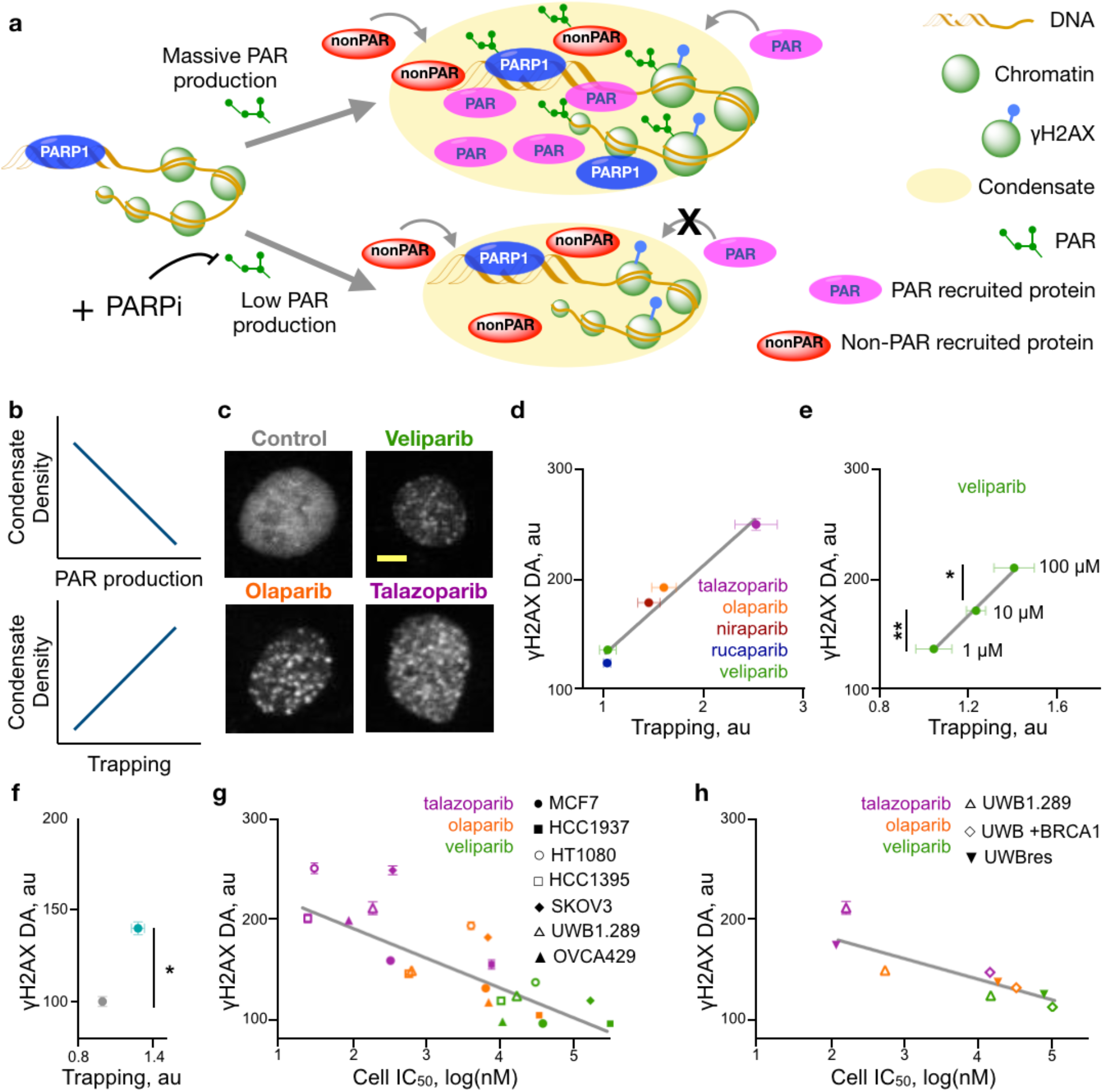
DNA damage biomolecular condensate density correlates to PARP trapping and cell line response to PARPi. (**a**) PAR is produced when PARP1 is activated upon binding DNA, which helps to establish a biomolecular condensate at the site of DNA damage by recruiting proteins. Other signaling also occurs, such as H2AX phosphorylation and protein recruitment in a PAR independent pathway. Under PARP inhibition, PAR-recruited proteins will be reduced and the relative density of other components will be higher. (**b**) Expected dependency of condensate density on PAR production and correlation to PARP trapping. (**c**) Representative γH_2_AX immunofluorescence images of HT1080 cell nuclei treated with 1 μM PARPi or control. (**d**) ICS DA of γH_2_AX IF in HT1080 cells treated with 1 μM PARPi overnight versus PARP1 trapping, with linear fit (gray line), shown are average with SEM, trapping - n=5, DA - n >= 536 cells, 3 biological repeats. (**e**) ICS DA of γH_2_AX IF in HT1080 cells treated with veliparib versus PARP1 trapping, with linear fit (gray line), shown are average with SEM, trapping - n=5, DA – n >= 1165 cells, 3 biological repeats, * p = 7×10^−18^, **p = 4×10^−16^, Student’s t test. (**f**) ICS DA of γH_2_AX IF in HT1080 cells treated with 10 nM GMX1778 for 24 hours or control versus PARP1 trapping, shown are average with SEM, trapping - n=5, DA - n >= 1285 cells, 3 biological repeats, * p = 1×10^−18^, Student’s t test. (**g**) ICS DA of γH_2_AX IF labeled cells treated with 1 μM PARPi overnight versus cell line IC_50_, shown are average with SEM, n >= 536 cells, 3 biological repeats, with linear fit (gray line). (**h**) ICS DA of γH_2_AX IF labeled cells treated with 1 μM PARPi overnight versus cell line IC_50_ for 3 UWB1.289 cell lines, shown are average with SEM, n >= 539 cells, 3 biological repeats, with linear fit (gray line).

To validate our ICS results, we measured individual condensate density of fluorescently labeled 53BP1 using anisotropy to quantify homo-FRET. In this approach, lower anisotropy is produced by increased homo-FRET, which is driven by increased fluorophore density^35^. We used a truncated 53BP1, which binds to H4K20me2^36^, labeled with mApple^37^ (Fig. S7e). The anisotropy of 53BP1-mApple in biomolecular condensates in HCC1937 cells was significantly lower than outside of condensates (Fig. S7f), demonstrating the increased density of condensates. However, similar to ICS measurements, condensate anisotropy was dependent on the presence and type of PARPi. We found significantly lower condensate anisotropy compared to control when cells were treated with 1 μM olaparib or talazoparib, but not with 1 μM veliparib, with no significant intensity differences (Fig. S7g). Furthermore, treatment with GMX1778 overnight significantly lowered condensate anisotropy without impacting condensate intensity (Fig. S7h & i), while increased veliparib lowered condensate 53BP1-mApple intensity suggesting recruitment is diminished at high veliparib concentrations (Fig. S7j & k).

We then correlated γH_2_AX ICS DA to cell line PARPi response for 3 PARPi in 7 cell lines (Fig. 3g). All cell lines showed a PARPi dependent γH_2_AX DA that correlates with PARPi efficacy (Fig. S8a), contrary to the allosteric impact of PARP1 DNA affinity or enzymatic properties of PARPi (Fig. S8c). In HCC1937 cells, a poor responder to PARPi treatment, talazoparib results clustered with olaparib treatment for other cell lines, and olaparib results similarly clustered with veliparib treatment, suggesting that γH_2_AX DA correlates to PARPi efficacy independent of both cell line and PARPi species. Unlike DA, some treatment conditions showed no significant increase in γH_2_AX levels over control, yet for all cell lines talazoparib treatment increased γH_2_AX levels (Fig. S8b). γH_2_AX DA also correlated to cell response in isogenic cell lines UWB1.289, UWB1.289 +BRCA1 and UWB1.289 resistant to olaparib (Fig. 3h and Fig. S8d-f). Thus, we concluded that PARPi altered DNA damage condensates, measured through condensate density, broadly correlate to both PARP trapping and PARPi efficacy.

We next modeled all the PARP1 binding events that occur within a condensate to determine if altered condensates could lead to observed PARP trapping. In our model (Fig. 4a), PARP1, PARPi and damaged DNA undergo binding and release following first order binding kinetics. When DNA-bound PARP1 binds NAD^+^, it rapidly PARylates nearby proteins – a model simplification based on the fast PARP1 enzymatic activity^38,39^ and rapid release of PARP1 in the absence of PARPi^17,19^. Our model also incorporates PARP1 diffusion away from and into condensates, includes non-diffusible PARylation targets (histones), stochastically selects which protein is PARylated and stochastically recruits other DNA binding proteins (DDR) into the condensate in a PAR-dependent manner.

**Fig. 4.**
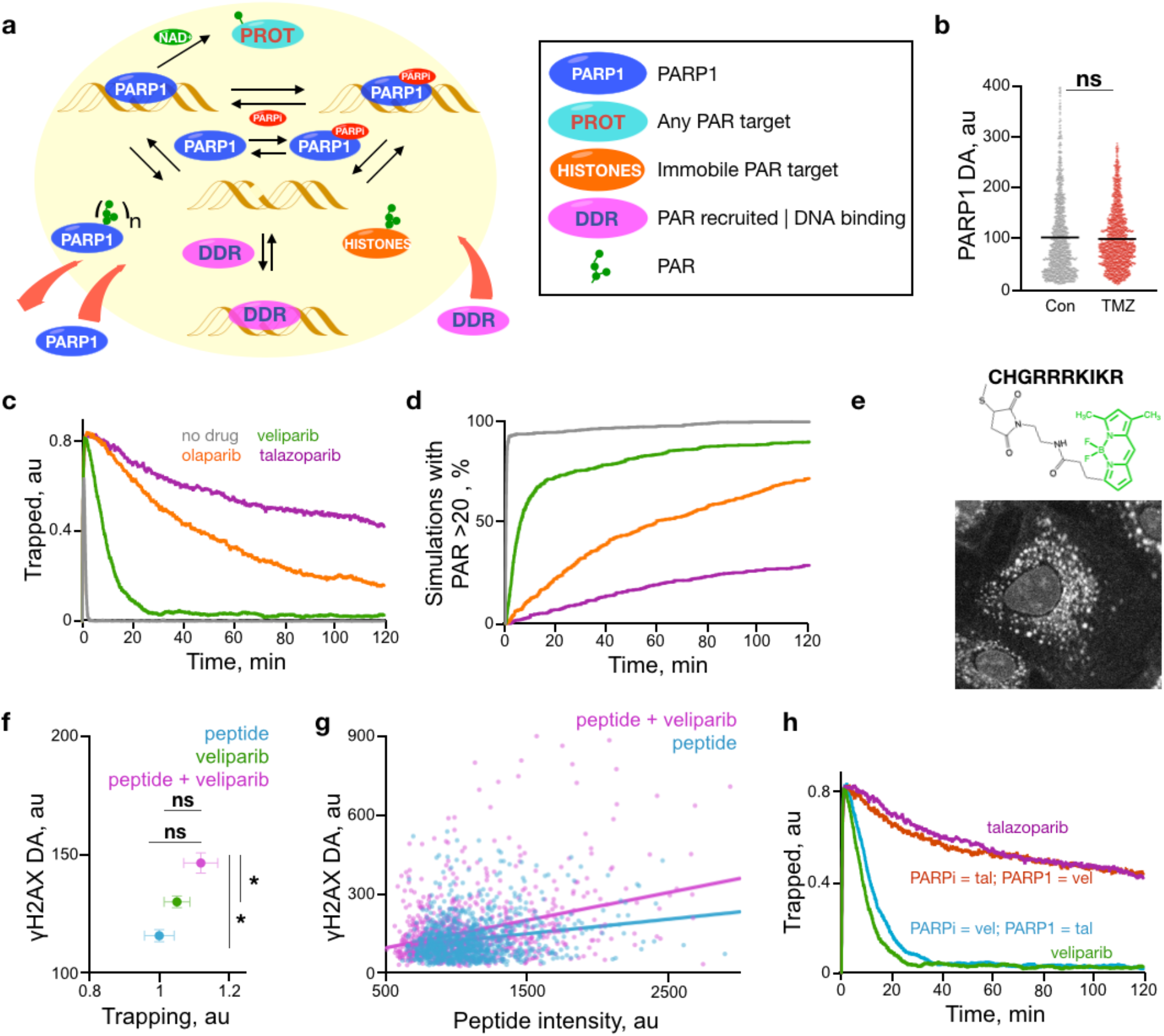
PARP trapping arises from altered PAR-dependent protein recruitment to biomolecular condensates. (**a**) Model of PARP1 activity during biomolecular condensate formation in response to DNA damage. PARP1, PARPi, DNA, and DDR (other DNA binding proteins in the DDR pathway) can bind to and dissociate from their targets. When DNA-bound PARP1 binds NAD^+^ a random protein is PARylated. PARylated PARP1 can exchange with unPARylated PARP1 outside the condensate and DDR proteins are recruited to condensates in a PAR-dependent manner. Histones represent PAR targets that cannot leave the condensate. (**b**) ICS DA of PARP1 IF in HT1080 cells treated overnight with 100 μM TMZ or control, shown are single cell values with average (black bar), n >= 1202 cells, 2 biological repeats. (**c-d**) Stochastic simulation results of (c) trapped PARP1 on DNA and (d) condensate PAR levels as a function of time in the absence or presence of 1 μM PARPi. (**e**) Representative image of 10 μM PAR binding peptide uptake in an HT1080 cell after 1 hour incubation. (**f**) ICS DA of γH_2_AX IF in HT1080 cells treated with 1 μM veliparib, 10 μM PAR binding peptide or the combination overnight versus PARP1 trapping, with linear fit (gray line), shown are average with SEM, trapping - n=5, DA - n >= 917 cells, 3 biological repeats, * p < 0.0005 (Student’s t test). (**g**) single nuclei ICS γH_2_AX DA of HT1080 cells versus PAR binding peptide nuclear intensity for cells treated with 10 μM PAR binding peptide (blue) or 10 μM PAR binding peptide plus 1 μM veliparib (pink) overnight, with linear fit, n >= 917 cells, 3 biological repeats. (**h**) Stochastic simulation results of trapped PARP1 on DNA as a function of time for veliparib (green), talazoparib (purple), an artificial PARPi with veliparib PARP1 binding constants and the talazoparib allosteric PARP1-DNA binding constants (blue), and an artificial PARPi with talazoparib PARP1 binding constants and the veliparib allosteric PARP1-DNA binding constants (red). All simulation data are an average of 1000 simulations.

We first sought to determine if PARP1 itself accumulates at sites of DNA damage. Experiments where DNA damage is induced through local, high intensity irradiation (UV laser exposure) show rapid accumulation of PARP1^40^. Yet, it remains uncertain if increased local PARP1 concentration occurs through multiple PARP1 proteins being recruited to single sites of DNA damage, or rather a high density of damaged DNA sites. When we induced DNA damage uniformly through incubation of cells with TMZ we found PARP1 does not aggregate (Fig. 4b), therefore the total PARP1 concentration in our model was kept constant. To implement our model we used the Gillespie algorithm^41^ (Fig. S9 and Supplemental Data). We tuned our simulation to achieve a trapped PARP duration in the absence of PARPi similar to observations in previous studies^17,19^ (Fig. S10). Here, PARP1 DNA occupancy was reduced to 20% within 2 minutes (Fig. 4c). Subsequently, talazoparib proved to be the most potent trapper, with a trapping half-life at 1 μM of 122 minutes compared to 44 and 6 minutes for olaparib and veliparib, respectively. We found trapping was associated with large decreases in PAR production and, as a result, PAR-binding protein recruitment (Fig. 4d and Fig. S10f-i).

Experimentally, inclusion of the PARG inhibitor PDD00017273 did not alter the veliparib or talazoparib induced γH_2_AX DA, but also did not impact HT1080 response to veliparib (Fig. S11a & b). However, a BODIPY FL labeled peptide containing the PAR-binding sequence of RNF146 (Fig. 4e) increased γH_2_AX DA in HT1080 cells, without increasing γH_2_AX levels (Fig. S11c), and significantly increased veliparib induced γH_2_AX DA with a correlation to non-significant increases in chromatin trapping (Fig. 4f). Furthermore, individual HT1080 cell γH_2_AX DA positively correlated to nuclear peptide accumulation (Fig. S11d), which was enhanced in the presence of veliparib (Fig. 4g). Our model and simulations (Fig. S10) suggest this PAR-binding peptide enhances veliparib trapping through interacting with PAR and preventing PAR-based recruitment. These results are similar to recent findings that limited PAR production through HPF1 knockout sensitizes cells to PARPi^42^.

We then ran simulations with reduced NAD^+^ concentrations and found increased trapping in a dose-dependent manner through limited protein recruitment (Fig. S12a & b). Simulations also showed veliparib concentration dependent trapping (Fig. S12c & d) similar to experimental chromatin trapping measurements. To determine which PARPi property (enzymatic inhibition of PAR production or allosteric induced PARP1 affinity for DNA) drives trapping, we ran simulations with synthetic PARPi containing a combination of veliparib and talazoparib binding constants (Fig. 4j). Here, the PARPi binding constants (enzymatic inhibition) proved to be the key regulator of PARP trapping, while allosteric changes in PARP1 affinity for DNA showed little impact (Fig. S12e and f). Simulations also suggested trapping through enzymatic inhibition was driven mainly by the k_off_ constant (Fig. S12g-j). Although clinical PARPi have comparable affinity for PARP1, their first order dissociation rate constants (k_off_) are strikingly different (Table S1), a potential explanation for why talazoparib is such potent trapper.

Next, we experimentally determined if PARPi-induced recruitment of DDR proteins correlates with response and focused on RPA1, a DNA single strand binding protein critical in various DDR pathways^43^. All 9 cell lines we measured showed increased RPA1 DA upon DNA damage with TMZ (Fig. 5a), demonstrating that RPA1 is recruited to DNA damage. But, the degree of recruitment did not correlate to cellular RPA1 expression. Surprisingly, when we measured RPA1 DA induced by PARPi treatment we found different recruitment trends across PARPi for each cell line (Fig. 5b). Here, cell lines more responsive to PARPi showed a decrease in RPA1 DA with increasing PARPi potency. Yet, for more resistant cell lines, RPA1 DA shows the opposite response (Fig. S13d). Remarkably, the relationship between RPA1 DA and PARPi IC_50_ for each cell line correlated to overall PARPi sensitivity (Fig. 5c). And, olaparib dose dependent RPA1 DA showed similar trends in HT1080 and HCC1937 cells (Fig. 5d), confirming that the degree of PARP inhibition is driving RPA1 recruitment. These results suggest resistant cell lines are able to recruit RPA1 in a PAR-independent manner while responsive cell lines lack this mechanism.

**Fig. 5.**
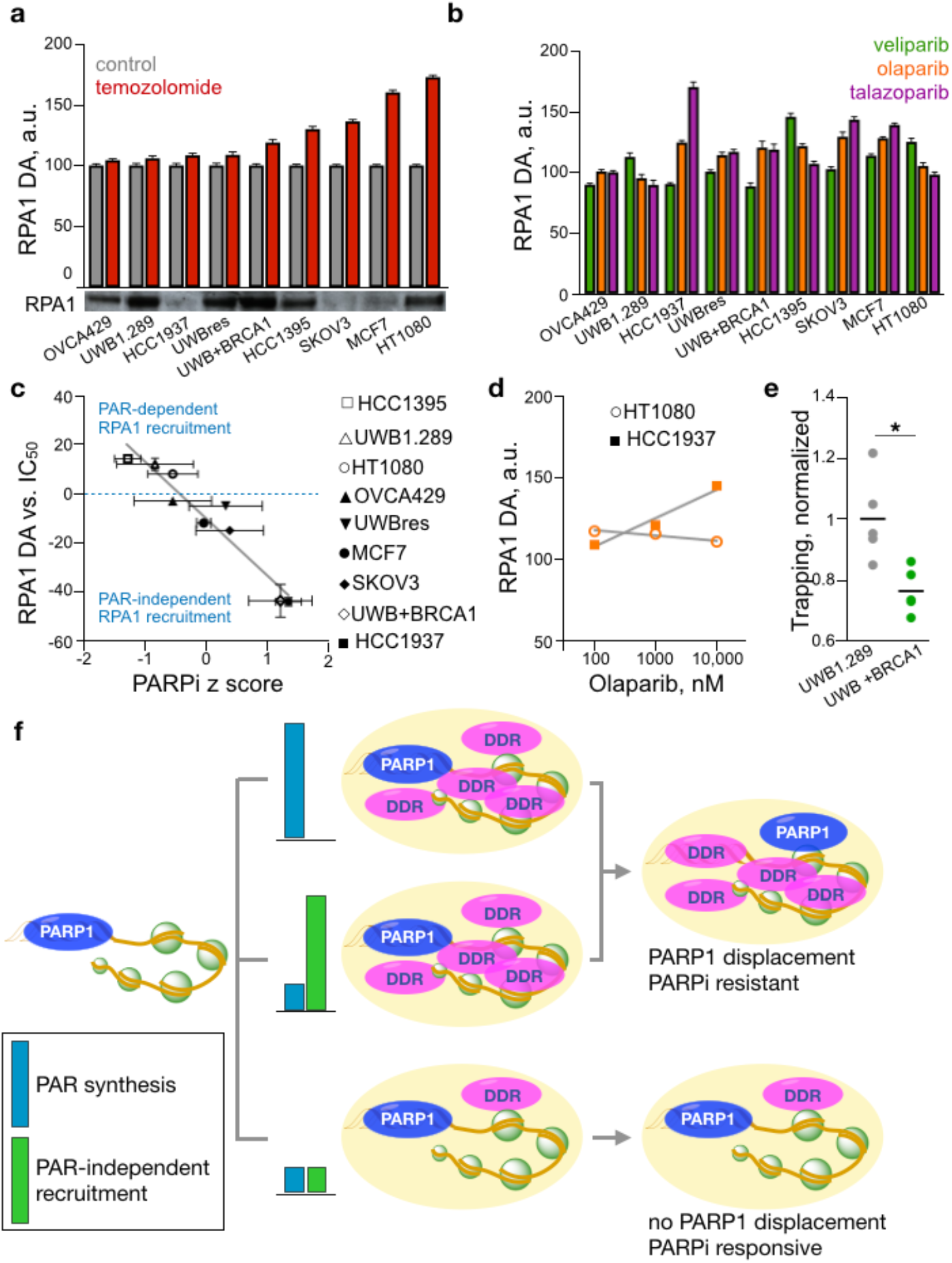
RPA1 recruitment correlates with cell sensitivity to PARPi. (**a**) RPA1 DA in untreated cells or cells treated with 100 μM TMZ overnight, shown are average with SEM, n >= 680 cells, 3 biological repeats, all cell lines except OVCA429 p < 0.05 for control vs. TMZ (Student’s t test). And, western blot RPA1 expression levels for each cell line. (**b**) RPA1 DA in cells treated with 1 μM PARPi overnight, shown are average, normalized to untreated control, with SEM, n >= 350 cells, 2 biological repeats. (**c**) The linear fit slope of RPA DA vs. cell line IC50 as a function of average PARPi z-score for each cell line. Shown are slope fit with SEM and average z-score over three PARPi with st. dev. (**d**) RPA1 DA as a function of olaparib concentration for HT1080 and HCC1937 cells, shown are average with SEM, n >= 504 cells, 3 biological repeats, and linear fit. (**e**) PARP1 trapping in UWB1.289 and UWB1.289 +BRCA1 cells treated with 1 μM talazoparib overnight, n=5, * p<0.05, Student’s t test. (**f**) Model of PARPi efficacy. DDR proteins are recruited by DNA damage condensates through PAR (blue bars) or PAR-independent mechanisms (green bars). The presence of DNA binding DDR proteins in the condensate results in increased competition for binding DNA which outcompetes PARP1. In the absence of DNA binding DDR proteins PARP1 remains “trapped” preventing progression of DNA damage repair.

Lastly, we found that PARP trapping was impacted by the presence of BRCA1. In isogenic UWB1.289 and UWB1.289 +BRCA1 cell lines, trapping was significantly higher in the absence of BRCA1 (Fig. 5e), suggesting that either BRCA1 reduces the number of DNA damage sites where PARP1 can be trapped or the presence of BRCA1 alters recruitment of proteins that compete with PARP1 in binding DNA. Our finding that RPA1 recruitment is inverted in these cells, but not γH_2_AX DA, by the presence of BRCA1 (Fig. 5c) indicates that recruitment does indeed play role in reduced trapping. These results agree with previous findings suggesting BRCA1 acts as a scaffold enabling protein and DNA interactions^44,45^. Yet, UWB1.289 cells resistant through culture in olaparib do not express BRCA1 (Fig. S8d) but have also gained the ability to recruit RPA1 in a PAR-independent manner. Therefore, our results suggest cell lines with the ability to recruit key DDR proteins in the absence PAR production are able to resist PARPi treatment (Fig. 5f). Broadly, the strong correlation of γH_2_AX density to IC_50_ and RPA1 recruitment to cell line PARPi sensitivity indicates condensate properties serve as a central mechanism governing response to PARP inhibition.

Our results suggest that, unlike with topoisomerase inhibitor trapping^46^, PARP trapping is not the physical stalling of PARP1 on DNA, but rather the persisting high fraction of damaged DNA bound by PARP1 in the absence of PAR-recruited competitive binding. PARP1 is one of the first proteins to bind damaged DNA not because it has the highest affinity (other proteins are known to have higher affinity^47,48^), but because of its high abundance^49^ and unique ability to locate damaged DNA^50^. When bound to DNA, PARP1 activity increases the local concentration of DNA binding proteins^30-32^ within growing condensates, altering the likelihood of PARP1 binding vacant DNA through increased competition. Indeed, absence of or mutations in DNA binding proteins that are recruited to DNA damage, such as XRCC1, produce hyperactive PARP, but also increase the PARP1-DNA interaction duration^19,51,52^. Thus, in the absence of DDR protein recruitment, once PARP1 dissociates from DNA it maintains a binding advantage to rebind DNA. Ultimately, all PARP inhibitors trap (Fig. 1a) and the PARPi dissociation constant is the essential characteristic controlling PARP trapping (Fig. 4j & Fig. S11).

## Methods

### Materials

All chemicals were purchased from Sigma unless otherwise noted. All drugs were purchased from Selleck Chemicals.

### Cell Culture

Cells were cultured in media supplemented with 10% fetal bovine serum and 1% penicillin/streptomycin. OVCA429 were originally obtained from Dr. Michael Birrer and cultured in DMEM. All other cell lines were obtained from ATCC and cultured in either RPMI or DMEM. PARP1 knockout cells were created using the LentiCrisprV2 system (a gift from Feng Zhang (Addgene plasmid # 52961))^53^ and the gRNA CGATGCCTATTACTGCACTG. HCC1937 expressing 53BP1-mApple were described previously^37^. UWB1.289 were grown to be olaparib resistant by culturing cells in increasing concentrations of olaparib up to 10 μM.

### Anisotropy and Confocal Microscopy

All fluorescence imaging was performed on an Olympus FV1000 multiphoton/confocal microscope. Anisotropy measurements were performed through custom modifications to the imaging system as previously described^22^. Briefly, linear polarization of two photon excitation light at 910 nm (MaiTai femtosecond laser, Spectra Physics) was controlled by a glan-thompson prism and half wave plate. Excitation was collected in orthogonal orientations through two detectors, parallel and perpendicular to excitation polarization, through a polarizing beam splitter that replaced a dichroic mirror in the emission filter cube. The alignment and detector gain and noise were validated each imaging session through measurements of a standard slide. Images were taken with an Olympus 25x XLPlan N objective, NA 1.05, and 3x digital zoom. Confocal imaging was performed with an Olympus XLUMPlanFL N 20x objective, NA 1.00 with chromatic correction, and 3x digital zoom.

### Drug Dissociation Assay

Cells were grown on 25 mm round, uncoated, sterilized glass coverslips in 6-well plates for 24 hours. Cells were then incubated with PARPi (1 μM) and/or labeled drug (500 nM) in imaging media (phenol-red free DMEM supplemented with 10% FBS and 1% pen-strep) for at least 15 minutes. Coverslips were removed from the 6-well plates, mounted onto a perfusion chamber (Warner Instruments), sealed with vacuum grease and perfused with a custom tubing setup. The chamber was then mounted onto the microscope and temperature was maintained at 37°C using a heating pad and feedback loop. Cells were initially located using fluorescent drug or autofluorescence and allowed to temperature equilibrate on the microscope stage for at least 5 minutes. Images were acquired using the anisotropy configuration as described above. After initial images were acquired, the chamber was perfused with 10x volume of the wash media containing fluorescent drug while chamber media and excess wash media were aspirated out of the chamber with vacuum. Images were then acquired at the desired time points while correcting for any drift. Association of fluorescent drug measurements were performed through rapid addition of media containing fluorescent drug (500 nM) and immediate image acquisition.

Nuclei were segmented in MATLAB and the average nuclear intensity and anisotropy were calculated using the *regionprops* function. The ΔrINT value was calculated as previously described^21^ (Supplemental Data). Values were then plotted as a function of time, with t=0 when PARPi was removed, in Prism (Graphpad) and a one phase association curve was fit to obtain the binding rate. The fitted association rate was used to determine the dissociation or association constant.

### Immunofluorescence

Cells were plated in 8-well 12-well slides (Thermo or Ibidi) to achieve 50% confluence overnight. Cells were then treated and fixed with 4% paraformaldehyde in PBS at room temperature for 10 minutes. Antibody staining was performed according to the manufacturers protocol. Following fluorescent secondary antibody labeling, prolong gold with DAPI (Cell Signaling Technologies) was added and samples were stored for up to 1 week prior to imaging.

### Image Correlation Spectroscopy

ICS analysis was performed in MATLAB as previously described^33^. Briefly, nuclei were segmented from the DAPI channel with the MATLAB function, *imbinarize*, converting the greyscale image to a binary dependent on an automatically defined threshold. The MATLAB function, *regionprops*, was used to identify single nuclei as objects. Each nucleus was used as a mask to perform ICS analysis on the antibody-stained channel, within size limitations for a nucleus (objects > 400 and < 3000 pixels). To ensure the analysis of antibody stained nuclei, nuclei under an average intensity of 200 au after background subtraction were excluded. The spatial autocorrelation functions were calculated using Fourier methods, then fit to a 2D Gaussian using a nonlinear least-squares algorithm^33^. Using the outputs of the fitting, the mean number of independent emitting fluorescent entities per focal spot area and subsequently the DA were determined^54^. DA measurements across experiments were normalized by the control in each dataset. The confocal imaging settings were kept the same across each dataset to provide comparable measurements per dataset. Any DA values exceeding 3 standard deviations from the mean were defined as incorrectly segmented and removed. Data were plotted in Prism and analysis performed in Prism or Excel.

### Condensate Anisotropy Measurements

Cells were grown on 12 mm round, uncoated, sterilized glass coverslips in 24-well plates for 24 hours. Cells were treated with PARPi and 100 μM TMZ for 1 hour or 10 nM GMX1778 overnight followed by 100 μM TMZ for 1 hour. TMZ was added to stimulate new DNA damage induced condensates in the presence of PARPi or NAD^+^ depletion. Cells were then transferred to the microscope for anisotropy imaging. Images were analyzed in MATLAB. Nuclei were segmented through intensity thresholding and condensates were segmented through further intensity thresholding with a higher threshold. Values of segmented nuclei and condensates were determined through the *regionprops* function. Segmentation was corrected through area restrictions to ensure that regions of interest were isolated condensates. The thresholding parameters were consistent throughout experiments. The average intensity and anisotropy of each segmented nuclei or condensate was then calculated. Data were plotted in Prism and analysis performed in Prism or Excel.

### Dose Response

Cells were plated in a 96-well plate at 2×10^3^ cells per well and treated with drug for 5 days in three replicate wells. Cell viability was then determined using Presto Blue (Thermo Fisher). Signal was averaged over 3 replicate wells and normalized to blank (no cells) as well as to the levels of untreated wells. A sigmoidal response curve was fit to the average of two experiments in Prism. PARPi z score was calculated for each PARPi using 9 cell lines. The average z score over three PARPi (veliparib, olaparib, talazoparib) was then determined as a representation of cell line PARPi sensitivity.

### Western Blotting

Cells were lysed with RIPA buffer (Thermo Fisher) containing protease inhibitors (Thermo Fisher), vigorously vortexed and incubated on ice for 15 minutes. Lysate was then pelleted by centrifugation at 14,000g for 15 min at 4 °C. Supernatant was transferred to a clean tube and total protein was determined using a BCA assay (Pierce). Protein was loaded into a NuPAGE gel (Thermo Fisher) and transferred to nitrocellulose paper (Thermo Fisher). Blocking and antibody labeling was then carried out according to the antibody manufacturer’s protocol. For PAR measurements, cells were treated with TMZ for 1 hour, or 1 μM H_2_O_2_ for 10 minutes in 6-well plates. Cells were trypsinized and washed once with PBS prior to cell lysis. Western blot expression levels were quantified by inverting the red channel of an image of the blot in FIJI, subtracting the background intensity and measuring the total intensity of the lane. These values where then normalized to the control signal.

### Chromatin Fraction Analysis

Cells in 6-well plates were treated with PARPi or GMX1778 for 24 hours. The media was removed, and cells were scraped off the culture dish in 1 ml PBS. The cells were pelleted by centrifugation at 500g for 4 minutes at 4°C and the solution was removed. Pellets were resuspended in 250 μl lysis buffer (10 mM HEPES, 10 mM KCl, 1.5 mM MgCl_2_, 340 mM sucrose, 10% glycerol, 0.1% Triton X-100, pH 7.9, protease inhibitors (Thermo Fisher)) and incubated on ice for 10 minutes. Nuclei were separated by centrifugation at 1300g for 5 minutes at 4°C and the solution was collected (cytoplasm fraction). Nuclei pellets were then washed with 100 μl lysis buffer lacking triton X-100 and centrifuged at 1300g for 2.5 minutes at 4°C. Pellets were then resuspended in 100 μl nuclear lysis buffer (50 mM HEPES, 250 mM KCl, 2.5 mM MgCl_2_, 0.1% Triton X-100, pH 7.5, protease inhibitors (Thermo Fisher)), vortexed at the highest setting for 5 seconds and incubated on ice for 10 minutes. The chromatin fraction was then pelleted by centrifugation at 15,000g for 10 minutes at 4°C and the solution was collected (nuclear fraction), the pellet was resuspended in 100 μl nuclear lysis buffer and re-pelleted by centrifugation at 15,000g for 10 minutes at 4°C. The pellet was then resuspended in 50 μl DNA release buffer (50 mM HEPES, 150 mM KCl, 2.5 mM MgCl_2_, 5 mM CaCl_2_, 0.05% Triton X-100, pH 7.5, protease inhibitors (Thermo Fisher)) and incubated at 37°C for 10 minutes. The solution was then centrifuged at 15,000g for 10 minutes at 4°C and the solution collected (trapped fraction). All solutes were collected for western blot analysis.

### PAR binding peptide synthesis

A peptide with the sequence CHGRRRKIKR was synthesized by Genscript. Peptide was dissolved in PBS containing 100mM molar tris-carboxyethylphosphine (TCEP), pH 7.4 to a final concentration of 1 mM in 3 ml and degassed with vacuum. BODIPY FL maleimide (Thermo) was dissolved in DMSO to a concentration of 60 mM in 200 μl. The dye was added to the peptide, flushed with argon and allowed to react overnight at 4°C. BODIPY FL peptide was purified by prep-HPLC and validated by LC-MS (Agilent). The fluorescent peptide was lyophilized over two days and dissolved in PBS at 1 mM.

## Author contributions

J.M.D. conceived the project and wrote the paper with editorial contributions from all authors; B.F. and J.M.D. performed experiments; A.A.G. and J.M.D. analyzed data; J.D. and J.M.D. developed and ran simulations. and R.W. and J.M.D. supervised the project.

## Data and materials availability

MATLAB files used to run models and simulations are available at: github.com/dubachLab/parpTrapping/.

## References

1. Kraus WL, Lis JT, PARP goes transcription. Cell 113, 677–683 (2003). 10.1016/s0092-8674(03)00433-1

2. Pascal JM, The comings and goings of PARP-1 in response to DNA damage. DNA Repair (Amst) 71, 177–182 (2018). 10.1016/j.dnarep.2018.08.022

3. Langelier MF, Planck JL, Roy S, Pascal JM, Structural basis for DNA damage-dependent poly(ADP-ribosyl)ation by human PARP-1. Science 336, 728–732 (2012). 10.1126/science.1216338

4. Alvarez-Gonzalez R, Jacobson MK, Characterization of polymers of adenosine diphosphate ribose generated in vitro and in vivo. Biochemistry 26, 3218–3224 (1987). 10.1021/bi00385a042

5. Ayyappan V, et al., ADPriboDB 2.0: an updated database of ADP-ribosylated proteins. Nucleic Acids Res 49, D261–D265 (2021). 10.1093/nar/gkaa941

6. Larsen SC, Hendriks IA, Lyon D, Jensen LJ, Nielsen ML, Systems-wide Analysis of Serine ADP-Ribosylation Reveals Widespread Occurrence and Site-Specific Overlap with Phosphorylation. Cell Rep 24, 2493–2505.e4 (2018). 10.1016/j.celrep.2018.07.083

7. Leung AKL, Poly(ADP-ribose): A Dynamic Trigger for Biomolecular Condensate Formation. Trends Cell Biol 30, 370–383 (2020). 10.1016/j.tcb.2020.02.002

8. Alemasova EE, Lavrik OI, Poly(ADP-ribosyl)ation by PARP1: reaction mechanism and regulatory proteins. Nucleic Acids Res 47, 3811–3827 (2019). 10.1093/nar/gkz120

9. Lord CJ, Ashworth A, PARP inhibitors: Synthetic lethality in the clinic. Science 355, 1152–1158 (2017). 10.1126/science.aam7344

10. Farmer H, et al., Targeting the DNA repair defect in BRCA mutant cells as a therapeutic strategy. Nature 434, 917–921 (2005). 10.1038/nature03445

11. Bryant HE, et al., Specific killing of BRCA2-deficient tumours with inhibitors of poly(ADP-ribose) polymerase. Nature 434, 913–917 (2005). 10.1038/nature03443

12. Murai J, et al., Trapping of PARP1 and PARP2 by Clinical PARP Inhibitors. Cancer Res 72, 5588–5599 (2012). 10.1158/0008-5472.CAN-12-2753

13. Murai J, et al., Stereospecific PARP trapping by BMN 673 and comparison with olaparib and rucaparib. Mol Cancer Ther 13, 433–443 (2014). 10.1158/1535-7163.MCT-13-0803

14. Ray Chaudhuri, A. et al., Replication fork stability confers chemoresistance in BRCA-deficient cells., Nature 535, 382–387 (2016). 10.1038/nature18325.

15. Helleday, T., The underlying mechanism for the PARP and BRCA synthetic lethality: clearing up the misunderstandings., Mol Oncol 5, 387–393 (2011). 10.1016/j.molonc.2011.07.001

16. Pommier Y, O’Connor MJ, de Bono J, Laying a trap to kill cancer cells: PARP inhibitors and their mechanisms of action. Sci Transl Med 8, 362ps17 (2016). 10.1126/scitranslmed.aaf9246

17. Zandarashvili L, et al., Structural basis for allosteric PARP-1 retention on DNA breaks. Science 368, (2020). 10.1126/science.aax6367

18. Hopkins TA, et al. Mechanistic Dissection of PARP1 Trapping and the Impact on In Vivo Tolerability and Efficacy of PARP Inhibitors. Mol Cancer Res 13, 1465–1477 (2015). 10.1158/1541-7786.MCR-15-0191-T

19. Shao Z, et al., Clinical PARP inhibitors do not abrogate PARP1 exchange at DNA damage sites in vivo. Nucleic Acids Res 48, 9694–9709 (2020). 10.1093/nar/gkaa718

20. Suskiewicz MJ, et al., HPF1 completes the PARP active site for DNA damage-induced ADP-ribosylation. Nature 579, 598–602 (2020). 10.1038/s41586-020-2013-6

21. Dubach JM, et al., Quantitating drug-target engagement in single cells in vitro and in vivo. Nat Chem Biol 13, 168–173 (2017). 10.1038/nchembio.2248

22. Dubach JM, et al., In vivo imaging of specific drug-target binding at subcellular resolution. Nat Commun 5, 3946 (2014). 10.1038/ncomms4946

23. Fu, D., Calvo, J. A., Samson, L. D., Balancing repair and tolerance of DNA damage caused by alkylating agents., Nat Rev Cancer 12, 104–120 (2012). 10.1038/nrc3185

24. Pessina F, et al., Functional transcription promoters at DNA double-strand breaks mediate RNA-driven phase separation of damage-response factors. Nat Cell Biol 21, 1286–1299 (2019). 10.1038/s41556-019-0392-4

25. Singatulina AS, et al., PARP-1 Activation Directs FUS to DNA Damage Sites to Form PARG-Reversible Compartments Enriched in Damaged DNA. Cell Rep 27, 1809–1821.e5 (2019). 10.1016/j.celrep.2019.04.031

26. Oshidari R et al., DNA repair by Rad52 liquid droplets. Nat Commun 11, 695 (2020). 10.1038/s41467-020-14546-z

27. Kilic S et al., Phase separation of 53BP1 determines liquid-like behavior of DNA repair compartments. EMBO J 38, e101379 (2019). 10.15252/embj.2018101379

28. Patel A, et al., A Liquid-to-Solid Phase Transition of the ALS Protein FUS Accelerated by Disease Mutation. Cell 162, 1066–1077 (2015). 10.1016/j.cell.2015.07.047

29. Altmeyer M, et al. Liquid demixing of intrinsically disordered proteins is seeded by poly(ADP-ribose). Nat Commun 6, 8088 (2015). 10.1038/ncomms9088

30. Nicolai S, et al., ZNF281 is recruited on DNA breaks to facilitate DNA repair by non-homologous end joining. Oncogene 39, 754–766 (2020). 10.1038/s41388-019-1028-7

31. Moore S, et al., The CHD6 chromatin remodeler is an oxidative DNA damage response factor. Nat Commun 10, 241 (2019). 10.1038/s41467-018-08111-y

32. Rona G, et al., PARP1-dependent recruitment of the FBXL10-RNF68-RNF2 ubiquitin ligase to sites of DNA damage controls H2A.Z loading. Elife 7, (2018). 10.7554/eLife.38771

33. Gopal AA, et al., Spatially Selective Dissection of Signal Transduction in Neurons Grown on Netrin-1 Printed Nanoarrays via Segmented Fluorescence Fluctuation Analysis. ACS Nano 11, 8131–8143 (2017). 10.1021/acsnano.7b03004

34. Natale F, et al., Identification of the elementary structural units of the DNA damage response. Nat Commun 8, 15760 (2017). 10.1038/ncomms15760

35. Varma R, Mayor S, GPI-anchored proteins are organized in submicron domains at the cell surface. Nature 394, 798–801 (1998). 10.1038/29563

36. Dimitrova N, Chen YC, Spector CL, de Lange T, 53BP1 promotes non-homologous end joining of telomeres by increasing chromatin mobility. Nature 456, 524–528 (2008). 10.1038/nature07433

37. Yang KS, Kohler RH, Landon M, Giedt R, Weissleder R, Single cell resolution in vivo imaging of DNA damage following PARP inhibition. Sci Rep 5, 10129 (2015). 10.1038/srep10129

38. Langelier MF, Ruhl DD, Planck JL, Kraus WL, Pascal JM, The Zn3 domain of human poly(ADP-ribose) polymerase-1 (PARP-1) functions in both DNA-dependent poly(ADP-ribose) synthesis activity and chromatin compaction. J Biol Chem 285, 18877–18887 (2010). 10.1074/jbc.M110.105668

39. Marsischky GT, Wilson BA, Collier RJ, Role of glutamic acid 988 of human poly-ADP-ribose polymerase in polymer formation. Evidence for active site similarities to the ADP-ribosylating toxins. J Biol Chem 270, 3247–3254 (1995). 10.1074/jbc.270.7.3247

40. Haince JF, et al., PARP1-dependent kinetics of recruitment of MRE11 and NBS1 proteins to multiple DNA damage sites. J Biol Chem 283, 1197–1208 (2008). 10.1074/jbc.M706734200

41. Gillespie DT, Exact stochastic simulation of coupled chemical reactions. The journal of physical chemistry 81, 2340–2361 (1977).

42. Prokhorova, E. et al., Serine-linked PARP1 auto-modification controls PARP inhibitor response., Nat Commun 12, 4055 (2021). 10.1038/s41467-021-24361-9

43. Dueva, R., Iliakis, G., Replication protein A: a multifunctional protein with roles in DNA replication, repair and beyond., NAR Cancer 2, zcaa022 (2020). 10.1093/narcan/zcaa022

44. Mark W, et al., Characterizatiion of segments from the central region of BRCA1: an intrinsically disordered scaffold for multiple protein-protein and protein-DNA interactions? J. Mol. Biol. 345, 275–287 (2005). 10.1016/j.jmb.2004.10.045

45. Foray N, et al. A subset of ATM-and ATR-dependent phosphorylation events requires the BRCA1 protein. EMBO J 22, 2860–2871. 10.1093/emboj/cdg274

46. Li TK, Liu LF, Tumor cell death induced by topoisomerase-targeting drugs. Annu Rev Pharmacol Toxicol 41, 53–77 (2001). 10.1146/annurev.pharmtox.41.1.53

47. Hey T, Lipps G, Krauss G, Binding of XPA and RPA to damaged DNA investigated by fluorescence anisotropy. Biochemistry 40, 2901–2910 (2001). 10.1021/bi002166i

48. Reardon JT, et al., Comparative analysis of binding of human damaged DNA-binding protein (XPE) and Escherichia coli damage recognition protein (UvrA) to the major ultraviolet photoproducts: T[c,s]T, T[t,s]T, T[6-4]T, and T[Dewar]T. J Biol Chem 268, 21301–21308 (1993).

49. Ludwig A, Behnke B, Holtlund J, Hilz H, Immunoquantitation and size determination of intrinsic poly(ADP-ribose) polymerase from acid precipitates. An analysis of the in vivo status in mammalian species and in lower eukaryotes. J Biol Chem 263, 6993–6999 (1988).

50. Rudolph J, Mahadevan J, Dyer P, Luger K, Poly(ADP-ribose) polymerase 1 searches DNA via a ‘monkey bar’ mechanism., Elife 7, (2018). 10.7554/eLife.37818

51. Hoch NC, et al., XRCC1 mutation is associated with PARP1 hyperactivation and cerebellar ataxia. Nature 541, 87–91 (2017). 10.1038/nature20790

52. Demin AA, et al., XRCC1 prevents toxic PARP1 trapping during DNA base excision repair. Mol Cell 81, 1–13 (2021). 10.1016/j.molcel.2021.05.009.

53. Sanjana NE, Shalem O, Zhang F, Improved vectors and genome-wide libraries for CRISPR screening. Nat Methods 11, 783–784 (2014). 10.1038/nmeth.3047

54. Wiseman PW, Petersen, NO Image correlation spectroscopy. II. Optimization for ultrasensitive detection of preexisting platelet-derived growth factor-beta receptor oligomers on intact cells. Biophys J 76, 963–977 (1999). 10.1016/S0006-3495(99)77260-7

55. Cambronne XA, et al., Biosensor reveals multiple sources for mitochondrial NAD+. Science 352, 1474–1477 (2016). 10.1126/science.aad5168

56. Fjeld CC, Birdsong WT, Goodman RH, Differential binding of NAD+ and NADH allows the transcriptional corepressor carboxyl-terminal binding protein to serve as a metabolic sensor. Proc Natl Acad Sci U S A 100, 9202–9207 (2003). 10.1073/pnas.1633591100

57. Langelier MF, Zandarashvili L, Aguiar PM, Black BE, Pascal JM, NAD<sup>+</sup> analog reveals PARP-1 substrate-blocking mechanism and allosteric communication from catalytic center to DNA-binding domains. Nat Commun 9, 844 (2018). 10.1038/s41467-018-03234-8

58. Shen Y, et al., BMN 673, a novel and highly potent PARP1/2 inhibitor for the treatment of human cancers with DNA repair deficiency. Clin Cancer Res 19, 5003–5015 (2013). 10.1158/1078-0432.CCR-13-1391

