## Supplemental Information for "Biomolecular condensates govern PARP inhibitor trapping and present mechanisms of resistance"

**Supplemental Data**

ODE Model of PARP-DNA binding

Our ODE model is solved in MATLAB using ODE45, a non-stiff solver. The script is available at github.com/dubachLab/parpTrapping/.

This model determines the duration of PARP1 engagement with DNA using previously established binding constants. Starting from the PARP1-DNA complex PARP1 can either release from DNA, self-PARylate to $PAR_n = 1$, or bind to PARPi to form the trapped complex. From trapped PARP (PARP1-DNA-PARPi complex) the PARPi-PARP1 pair can dissociate from DNA with the drug occupancy remaining intact or PARPi can dissociate from the PARP1-DNA complex. In our model, each PARylation of PARP1 creates a new species that can undergo any of the steps as the previous PARP1 species. Once PARP1 dissociates from DNA it cannot rebind, thus we are modeling the duration of a singular PARP1-DNA binding event. Here, our initial values consist of complete PARPi-PARP1-DNA complexation with no prior PARylation. For simplicity we consider self PARylation as the only PARylation reaction. Although PARP1 does PARylate numerous other proteins, they theoretically do not impact PARP1-DNA affinity. Thus, omission of these alternative PARylation reactions only impacts the number of PARylation events prior to PARP1 forced removal from DNA, or the effect of PAR accumulation on PARP1 affinity for DNA. However, as is demonstrated by the results, PARP1 dissociation from DNA primarily occurs prior to PARylation forced removal, therefore we do not consider the absence of other PARylation targets to impact the results. Adjusting the rate constants, and other reaction parameters, provides a route to investigate how rates impact

PARP1-DNA binding duration. These constants and parameters are defined and described below.

**SPECIES**:

PARP1: Poly(ADP-ribose) polymerase. Here, the PAR status of PARP1 defines the species. For example, unPARylated PARP1 is a different species than PARP1 with one ADP-ribose modification, which is a different species than PARP1 with two ADP-ribose groups.

PARPi: PARP inhibitors.

PAR: Poly(ADP-ribose). PAR can only exist PARP1.

**REACTIONS AND BINDING EVENTS**:

For each round of PARP1 PARylation the following reactions and binding events can occur.

- PARP1 dissociation from DNA.
- PARP1-DNA association with PARPi.
- PARPi dissociation from PARP1-DNA.
- PARPi-PARP1 dissociation from DNA.
- PARP1 self-PARylation.

**CONSTANTS AND VARIABLES**:

*Rounds*. The number of PARP1 self-PARylation events that occur prior to electrostatic removal of PARP1 from DNA. Here the default is 500. A higher number of rounds (greater PARylation prior to forced release) only slightly increased PARP1-DNA residence in the absence of drug without impacting trapping in the presence of 1 μM olaparib (Fig. S3b). However, a lower number of rounds decreased the PARP1-DNA residence and olaparib induced PARP1 trapping. The lower number of rounds allows PARP1 to dissociate much faster due to: 1) the $\gamma$ affinity correction factor being spread out over fewer rounds, and 2) complete PARylation ($PAR_n$ = rounds) forcing removal of PARP1 from DNA.

$k_a$ *(trap)*: The association constant of PARPi for PARP1. This rate constant has previously been shown to be independent of PARP1 engagement to DNA[18]. Values were taken as an average of previous measurements (Table S1). The default values used were: veliparib 1.8e6 $M^{-1}s^{-1}$; olaparib 2e5 $M^{-1}s^{-1}$; talazoparib 3.6e5 $M^{-1}s^{-1}$.

$k_d$ *(trap)*: The dissociation of PARPi from PARP1-DNA. This rate constant has previously been shown to be independent of PARP1 engagement to DNA[18]. Values were taken as an average of previous measurements (Table S1). The default values used were: veliparib 5e-3 $s^{-1}$; olaparib 3e-4 $s^{-1}$; talazoparib 8e-5 $s^{-1}$.

*Drug concentration*: The PARPi concentration in the cell.

*$NAD^+$ concentration*: The default concentration of $NAD^+$ in the nucleus was 100 μM[55,56]. The concentration of $NAD^+$ in the ODE solution greatly impacted the PARP1-DNA residency time in the absence of drug, while having slight impact of PARP1 trapping in the presence of 1 μM olaparib (Fig. S3c). A higher concentration increased release of PARP1, while a lower concentration extended PARP1-DNA binding. With olaparib present the increased trapping reaches a plateau at 10 μM, 10% of the actual nuclear concentration. However, in the absence of

drug the impact doesn't plateau until a concentration of 100 nM, where the PARP1-DNA residency is similar to that of the presence of 1 μM PARPi under normal $NAD^+$ concentrations. Thus, below 100 nM, PARP1 release from DNA is driven solely by PARP1 affinity for DNA and independent of the impact of PARylation.

$k_a$ *(par)*: The association constant of $NAD^+$ for PARP1. Here, for simplicity, self-PARylation occurs immediately upon $NAD^+$ binding PARP1-DNA. The default value was 5e5 $M^{-1}s^{-1}$, derived from previously measured $K_m$ values[37]. The impact of $k_a$ *(par)* is nearly identical to the concentration of $NAD^+$ since the rate of PARylation is dependent on the product of the two (Fig. S3d).

$\gamma$: PARP1 loses affinity for DNA as it becomes PARylated, which arises from electrostatic repulsion. Thus, $\gamma$ is a correction factor that confers PAR dependency on the affinity of PARP1 for DNA. This correction factor serves to increase the dissociation rate constant of both PARP1 and PARPi bound PARP1 for DNA. The value of $\gamma$ is the number of order of magnitudes that the rate constants will change over the number of rounds. The correction is applied in a log scale to the binding rate constants. The default of $\gamma$ is 2, meaning that the binding constants will change two orders of magnitude over the number of rounds. The value of $\gamma$ slightly altered the PARP1-DNA residence time in the absence of drug and PARP1 trapping in the presence of 1 μM olaparib (Fig. S3e). Lower values of $\gamma$ increase the PARP1-DNA interaction through maintaining the default PARP1 DNA and PARPi-PARP1 DNA affinity during increasing PARylation. Thus, when $\gamma = 0$, there is a two-phase response in the absence of drug - initially where PARP1 dissociates from DNA and a more rapid decrease where complete PARylation causes removal from DNA. Higher values of $\gamma$ decrease the PARP1-DNA interaction through a greater loss of DNA affinity as a function of PARylation.

*$k_d$ (rel)*: The dissociation constant of PARP1 from DNA. This rate is dependent on the correction factor $\gamma$ and the degree of PARP1 PARylation. The default value was 3.36e-3 $s^{-1}$ (Table S1). This constant had minimal impact on PARP1-DNA interactions (Fig. S3f). Lower values (greater affinity) did not impact PARP1 trapping in the presence of 1 μM olaparib, demonstrating that PARP1 is largely occupied by olaparib. Higher values in the presence of 1 μM olaparib lowered the initial starting position as PARP1 dissociated from DNA prior to binding olaparib. In the absence of drug, higher values caused more rapid dissociation from DNA, as expected, while lower values shifted dissociation of PARP1 from DNA toward completely PARylated forced release, creating a delayed, steeper response.

*$k_d$ (rel PARPi)*: The dissociation constant of PARPi-PARP1 from DNA. This rate is dependent on the correction factor $\gamma$ and the degree of PARP1 PARylation. The default values used were (Table S1): veliparib 4.08e-3 $s^{-1}$; olaparib 2.31e-3 $s^{-1}$; talazoparib 2.52e-3 $s^{-1}$.

Apparent $k_{off}$ measurement model

Here we developed an assay to measure the intracellular dissociation constant of PARPi through binding of fluorescently labeled olaparib (Fig. 2a). PARPi at 1 μM is first applied to cells to saturate the target. Fluorescent olaparib at 500 nM is the added and allowed to diffuse into the cell to establish a concentration equilibrium. The higher concentration and affinity of the clinical

PARPi ensures the target remains occupied. The unbound clinical PARPi is then removed through excess washing while the fluorescent olaparib remains. The competition experiment to measure intracellular observed dissociation constants is set up so that t=0 occurs once free, unlabeled drug is removed from the system. The dynamic changes in drug target occupancy can then be defined by a set of ordinary differential equations:

$$\frac{d[RD]}{dt} = -k_{off}[RD] + k_{on}[D][R] \tag{1}$$

$$\frac{d[R]}{dt} = k_{off}[RD] + k_{off}[RD_f] - k_{on}[D_f][R] - k_{on}[D][R] \tag{2}$$

$$\frac{d[RD_f]}{dt} = -k_{off}[RD_f] + k_{on}[D_f][R] \tag{3}$$

where, [R] = the concentration of free target, [D] = the concentration of free drug, $[D_f]$ = the concentration of fluorescently labeled drug, [RD] = the concentration of target bound drug, and $[RD_f]$ = concentration of target bound fluorescently labeled drug. At t=0 all target is bound by drug ($[R]_{tot}$ = [RD]). We also assume that all free drug is removed from the system ($k_{on}[D][R] = 0$). Furthermore, due to the high concentration of fluorescently labeled drug, we assume that all target will be immediately bound by fluorescent drug once unlabeled drug dissociates and that, once engaged to fluorescent drug, target will remain occupied, or $k_{on}[D_f][R] >> k_{off}[RD_f]$. Thus, the amount of fluorescent drug bound to target is equal to the total amount of target minus the unlabeled drug bound target:

$$[RD_f] = [R]_{tot} - [RD] \tag{4}$$

Solving the differential equations with the above criteria we obtain:

$$[RD] = [R]_{tot} \exp\left(-k_{off}t\right) \quad (5)$$

$$\left[RD_f\right] = [R]_{tot}(1 - \exp\left(-k_{off}t\right)) \quad (6)$$

with the boundary conditions:

$$at\ t = 0, [RD] = [R]_{tot}$$

$$at\ t = \infty, [RD] = 0$$

$$at\ t = 0, \left[RD_f\right] = 0$$

$$at\ t = \infty, \left[RD_f\right] = [R]_{tot}$$

This ODE solution closely approximates discreet ODE solvers (Fig. S4a). This solution only slightly overestimates the amount of bound fluorescent drug as the clinical drug dissociates over time when compared to a stiff ODE solver (ODE15s, MATLAB). Here, we use a drug-target dissociation constant of 3 x $10^{-4}$ $s^{-1}$, a fluorescent drug-target dissociation constant of 3 x $10^{-6}$ $s^{-1}$, a fluorescent drug-target association constant of 2 x $10^{4}$ $M^{-1}s^{-1}$, and a fluorescent drug concentration of 500 nM. The difference between the ODE solution and solver largely arises from the presence of a small amount of free target calculated by the ODE solver (Fig. S4a). In our model we assume any free target is immediately occupied by fluorescent drug. However, both capture the association of fluorescent drug as clinical drug dissociates from the target. And, our approximation is valid through the expected range of dissociation rates (Fig. S4b)

Here we determine the amount of target bound fluorescent drug through measuring the fluorescence anisotropy in each imaging voxel[21,22]. Anisotropy is an ensemble average of discreet anisotropy states of all fluorophores in each voxel, therefore anisotropy can be defined as the following:

$$r = \sum_{n=1}^{i} \frac{r_n N_n}{N_{tot}} \tag{7}$$

where r = anisotropy, $r_n$ is the anisotropy of the fluorophore state n, $N_n$ is the number of fluorophores in the state n, and $N_{tot}$ is the total number of fluorophores in the voxel. Here, we assume a two-state system where the fluorescent drug can be labile in the cell or bound to the target. Therefore:

$$r = \frac{r_{bound}\left[RD_f\right] + r_{labile}[D_f]_{labile}}{[D_f]_{tot}} \tag{8}$$

where $r_{bound}$ = anisotropy of a target bound fluorescent drug, $[RD_f]$ = the concentration of target bound fluorescent drug, $r_{labile}$ = anisotropy of unbound fluorescent drug, $[D_f]_{labile}$ = the concentration of unbound fluorescent drug, and $[D_f]_{tot}$ = the total concentration of fluorescent drug. Additionally, the total concentration of drug in each voxel is equal to the sum of unbound and bound:

$$[D_f]_{tot} = [D_f]_{labile} + [RD_f] \tag{9}$$

Solving for bound fluorescent drug we derive the expression:

$$[RD_f] = \frac{r - r_{labile}}{r_{bound} - r_{labile}}[D_f]_{tot} \tag{10}$$

If we assume no fluorophore quenching or photobleaching and a linear relationship between fluorescence intensity and fluorescent drug concentration, we can express voxel intensity as:

$$Int = \gamma[D_f]_{tot} \tag{11}$$

where Int = measured intensity and γ is a constant. Combining equations 6, 10 and 11 we derive:

$$(r - r_{labile})Int = (r_{bound} - r_{labile})\gamma[R]_{tot}(1 - \exp(-k_{off}t)) \tag{12}$$

where ($r_{bound}$ - $r_{labile}$) γ $[R]_{tot}$ is a constant expression. Thus, for each voxel, the measured anisotropy corrected by the unbound anisotropy value and multiplied by the intensity is a function of time. This value, ($r$ – $r_{labile}$)Int, which we refer to as ΔrINT, is a measurement of the amount of fluorescent drug bound to target[20]. As such, it can be directly used to determine the dissociation constant ($k_{off}$) of unlabeled clinical drug in cells. Here, we determine $k_{off}$ by fitting an exponential using Prism to the ΔrINT signal over time following removal of free clinical drug from the system.

Stochastic Simulation Model

The Gillespie simulation is a stochastic algorithm to advance the binding reactions within a biomolecular condensate through Monte-Carlo inversion steps. Each reaction in the system is associated with a variable propensity function $a_n$, where n is the number of possible events. The propensity function is dependent on both the reaction/binding rate and the concentration of the species involved. Each propensity function is updated after every time step. The simulation begins at $t=t_0=0$, and two random numbers are initialized, $r_1$ and $r_2$. The first random number generates the time step, τ, otherwise referred to as the sojourn time, at which the next reaction occurs, through the following equation:

$$\tau = \frac{1}{\sum_n a_n} \ln\left(\frac{1}{r_1}\right) \tag{13}$$

The second random number selects the reaction that occurs at $t_1 = t_0+\tau$, by minimizing the index n such that:

$$\sum_{i=1}^{i} a_i > r_2 \sum_n a_n \tag{14}$$

Here, the reaction that satisfies the above criteria is selected and the distribution of molecules in the system is updated. The simulation then continues at $t = t_1 = t_0+\tau$, where $\tau$ and the selected reaction are determined through newly generated random numbers, and repeats until $t >= t_{end}$, or a maximum number of reactions has occurred.

In our model, if a PARylation reaction is selected, which can only occur if uninhibited PARP1 is bound to DNA, a random PARP1 protein or histone is selected based on the total number of each species in the system. Each PARP1 is separately identified based on the number of ADP-ribose molecules that have previously been attached, however histones serve as a pool of PAR targets and are not tracked individually. After PARylation, either the PAR status of the selected PARP1 is updated, or the histone PAR level is updated, depending on which was selected for PARylation, and the total system PAR level is updated.

If a PARP1 exchange reaction is selected, then the PARP1 that was randomly chosen will be replaced with a PARP1 that lacks ADP-ribosylation – an exchange with PARP1 from outside the condensate. Here, we omit exchange of unPARylated PARP1, since that would not impact the status of the system. Thus, PARP1 in the system that has no PARylation cannot be selected for PARP1 exchange. After an exchange the total PAR in the condensate is updated.

At each step a random number is generated to stochastically decide if a DNA binding protein will enter the condensate. The likelihood of this protein entering the condensate is a function of both the protein threshold parameter (*p.prot_thresh*) described below and the total PAR in the condensate. If a DNA binding protein enters the condensate the number of proteins is updated, here these proteins cannot leave the condensate.

We implemented the Gillespie algorithm in MATLAB using the species, reactions, and constants described below. The script is available at github.com/dubachLab/parpTrapping/.

**SPECIES**:

*Damaged DNA*: Each condensate contains a single site of DNA damage that is capable of binding PARP1, the PARPi-PARP1 complex or DDR.

*DDR*: DNA binding proteins that can be recruited to condensates in a PAR dependent manner. DDR can bind to damaged DNA.

*PROT*: Proteins that are the target of PARP1 PARylation - PARP1 or histones.

*Histones*: Histone proteins are PAR targets and have a fixed condensate concentration. Here, histones are representative of any PAR targets that accumulate in the condensate and do not leave, however they are unable to bind damaged DNA.

*PARP1*: Poly(ADP-ribose) polymerase. Here the PAR status of PARP1 defines the species. For example, unPARylated PARP1 is different than PARP1 with one ADP-ribose modification, which is different that PARP1 with two ADP-ribose groups, and so on.

*PARPi*: PARP inhibitors.

*PAR*: Poly(ADP-ribose). PAR can only exist on histones or PARP1.

**REACTIONS**:

- DDR binding to damaged DNA.
- DDR dissociating from damaged DNA.

*PARP1 reactions*. The PARylation state of PARP1 defines each PARP1 molecule and represents a different species. Therefore, the number of PARP1 reactions below is defined by the number of rounds, where each round of PARylated PARP1 (PAR = 0,1,2…rounds) can undergo the following reactions.

- PARP1 binding to DNA
- PARP1 dissociating from DNA
- PARP1 binding PARPi
- PARP1 dissociating from PARPi
- PARP1-PARPi binding to DNA
- PARP1-PARPi dissociating from DNA
- PARPi binding to PARP1-DNA
- PARPi dissociating from PARP1-DNA
- PARylated PARP1 exchanging with un-PARylated PARP1
- PARP1 binding to $NAD^+$ and PARylating a protein through stochastic selection.

**CONSTANTS AND VARIABLES**:

*iterations*: The number simulations to run and average.

*end_time*: the duration of the simulation in seconds.

*drug_concentration*: The concentration of PARPi in the cell, this is a constant.

*rounds*: The length of PAR polymer on PARP1 when PARP1 loses affinity for damaged DNA. This value also defines the number of PARP1 states that can exist where each PARP1 state (round) is the length of PAR polymer attached to PARP1. The default value is 50. Under normal conditions in the presence or absence of PARPi, PARP1 does not progress beyond round 50 (Fig. S9a). However, in the absence of PARP exchange, γ and histones, PARP1 can progress beyond 100 rounds (Fig. S9a).

*PARylation protein target (p.histones)*: PARP1 PARylates numerous proteins and prominently PARylates histones. The value *p.histones* defines a fixed concentration PAR receiving proteins in the condensate. This concentration does not change, and the length of PAR polymer is not limited. PARylation of histones when DNA bound PARP1 binds $NAD^+$ is a stochastic selection that competes with PARylation of PARP1 proteins, therefore the relative concentrations of histones and PARP1 govern the probability of PARylation for each species. Here we do not track the PARylation status of individual histones, rather upon histone PARylation the entire non-reversible PAR status of the condensate increases by one unit of PAR. Simulations demonstrate that the concentration of histones does not alter PARP trapping in the presence of 1 μM olaparib, however there is profound dependency on the PARP-DNA residency in the absence of PARPi (Fig. S10a). Therefore, the default value of *p.histones* was tuned to achieve a PARP1-DNA curve with a duration similar to experimental observations[17,19]. The default value is 10x the concentration of PARP1.

*Initial PARP1 concentration (p.PARP0)*: The concentration of PARP1 in the condensate. PARP1 concentration was approximated at 1 μM[49]. The concentration of total PARP1 in the condensate is constant. The default value is 5 molecules per damaged DNA site. The concentration of PARP1 only impacts trapping when the relative concentration compared to initial DNA binding protein is altered (Fig. S10b).

*Initial DNA binding protein concentration in the condensate (p.prot0)*: The initial concentration of DNA binding proteins in the condensate (DDR). DNA binding proteins are recruited to the condensate in a PAR dependent manner and modeled to not leave the condensate. The default value is 1, however half the simulations are run at the default value and the other half run at *p.Proto* = 0 to capture the stochasticity of protein distribution throughout the nucleus. Therefore, the average default concentration is 10% of the default PARP1 concentration. The initial PARP1:DNA binding protein ratio impacts the maximum amount of PARP1 that is initially trapped in the presence of 1 μM olaparib, but does not alter the rate of loss of trapped PARP1 (Fig. S10b).

*PAR dependent PARP1 DNA affinity (p.gamma)*: PARP1 loses affinity for DNA as it becomes PARylated, arising from electrostatic repulsion of negatively charged PAR and DNA. Thus, $\gamma$ is a correction factor that confers PAR dependency on the affinity of PARP1 for DNA. This correction factor serves to both lower the association rate constant and increase the dissociation rate constant of both PARP1 and PARPi-bound PARP1 for DNA. The value of $\gamma$ is the number of orders of magnitude that the rate constants will change over the number of rounds. The correction is applied in a log scale to the binding rate constants. The default of $\gamma$ is 2, meaning that the binding constants will change two orders of magnitude over the number of rounds. This constant had no impact on PARP1 trapping in the presence of olaparib, however it did impact DNA residence time in the absence of drug (Fig. S10c) Therefore, the value was tuned to achieve the previously observed PARP1 DNA residence time in cells[17,19].

*PARP exchange (p.exchange)*: Shao, et al., demonstrated fluorescent PARP1 recovery after photobleach at sites of UV-induced DNA damage with an initial rate of approximately 20% recovery over 2 seconds[19]. We interpreted this initial rate to represent exchange of PARylated PARP1 for unmodified PARP1 at the site of DNA damage. Therefore, 10% of PARP1 molecules in our system exchange with the surrounding PARP1 every second. We fit our exchange rate constant to achieve this approximate rate and derived the expression *p.exchange* = 0.2/*p.PARP0*, where *p.PARP0* is the amount of PARP in the system. However, the PARP1 exchange rate had no impact on olaparib induced PARP1 trapping nor PARP1 DNA residency in the absence of drug (Fig. S10d).

*PAR dependent DNA binding protein recruitment (p.prot_thresh)*: DNA binding proteins are recruited into the condensate as a function of PAR accumulation. The variable *p.prot_thresh* defines the relationship between PAR and DNA binding protein recruitment. Stochastic selection of protein recruitment is dependent on the value of *p.prot_thresh*, with a higher value requiring greater PAR accumulation for each protein recruited. The default value of 2 was tuned to achieve the previously observed PARP1 DNA residence time in the absence of PARPi in cells[17,19]. This constant impacts PARP1 trapping in the presence of olaparib (Fig. S10e), but not PARP1 DNA association in the absence of PARPi. The impact of this constant demonstrates the trapping dependency of DNA binding protein (DDR) recruitment.

*DNA in the condensate (p.DNA0)*: Our simulation models a single section of damaged DNA, therefore this value is set to 1 and is constant.

*$NAD^+$ concentration (p.NAD)*: The default concentration is 100 μM based on previous measurements[55,56].

*Dissociation of PARPi from PARP1 (p.kd_trap)*: The dissociation constant of each PARPi from PARP1. Values were taken as an average of previous measurements (Table S1). The default values used were: veliparib 5e-3 $s^{-1}$; olaparib 3e-4 $s^{-1}$; talazoparib 8e-5 $s^{-1}$. This constant impacted PARP1 trapping when the values for olaparib were artificially adjusted (Supplemental Data Fig. 12e). Lower values (higher affinity) increased trapping through decreasing the dissociation of PARPi from PARP1, which lowers the amount of PAR generated.

*Association of PARPi to PARP1 (p.ka_trap)*: The association constant of each PARPi on PARP1. Values were taken as an average of previous measurements (Table S1). Rates were corrected to account for the interpretation of PARP1 concentration relative to DNA through multiplying the actual rate by $10^{-7}$. This rate correction was performed for all association constants. The default values used were: veliparib 1.8e-1 $M^{-1}s^{-1}$; olaparib 2e-2 $M^{-1}s^{-1}$; talazoparib 3.6e-2 $M^{-1}s^{-1}$.

*Association of $NAD^+$ on PARP1 (p.ka_PAR)*: The binding association constant of $NAD^+$ to PARP1. For simplicity, the PARylation reaction was modeled as instantaneous upon PARP1 binding to $NAD^+$. This binding constant was defined by previous measurements of PARP1 $K_M$ when bound to damaged DNA[38,39] and the affinity of an $NAD^+$ analog[57]. The default value, corrected for PARP/DNA concentration interpretation, was 5e-2 $M^{-1}s^{-1}$.

*Association of DNA binding protein to DNA (p.ka_prot)*: The binding association constant of DNA binding proteins to DNA was defined by the PARP1-DNA association constant. This value was set to 1e+5 $M^{-1}s^{-1}$. Corrected for PARP1/DNA concentration interpretation, the default was 1e-2 $M^{-1}s^{-1}$. This value was set to be the same as the association constant of PARP1 binding to DNA.

*Dissociation of DNA binding protein from DNA (p.kd_prot)*: The dissociation constant of DNA binding proteins from DNA was defined as an order of magnitude lower than the PARP1-DNA dissociation constant. This value was chosen to capture progression along the DNA repair pathway once DDR DNA binding proteins bind to damaged DNA. The default value was 4e-4 $s^{-1}$.

*Dissociation of PARP1 from DNA (p.kd_rel0)*: The dissociation constant of PARP1 from DNA. The value is an average of previous measurements (Table S1). This rate is dependent on the correction factor $\gamma$ and, thus, the degree of PARP1 PARylation. The default value was 3.36e-3 $s^{-1}$.

*Association of PARP1 to DNA (p.ka_rel0)*: The association constant of PARP1 on DNA. Because of the large variation in values between previous measurements the value is an average of previous measurements (Table S1) and set to 1e+5 $M^{-1}s^{-1}$. This rate is dependent on the correction factor $\gamma$ and, thus, the degree of PARP1 PARylation. Rates were corrected to account for the interpretation of PARP1 concentration relative to DNA through multiplying the actual rate by $10^{-7}$. The default was 1e-2 $M^{-1}s^{-1}$.

*Dissociation of PARPi-PARP1 from DNA (p.kd_relD0)*: The dissociation constant of PARPi bound PARP1 from DNA. Values were taken from previous findings[15]. This rate is dependent on the correction factor $\gamma$ and, thus, the degree of PARP1 PARylation. The default values used were: veliparib 4.08e-3 $s^{-1}$; olaparib 2.31e-3 $s^{-1}$; talazoparib 2.52e-3 $s^{-1}$. This constant impacted PARP1 trapping when the values for olaparib were artificially adjusted (Fig. S11f). Lower values (higher affinity) increased trapping through changing the competitive binding probabilities of PARP1 relative to DDR proteins.

*Association of PARPi-PARP1 to DNA (p.ka_relD0)*: The association constant of PARPi-bound PARP1 on DNA. Because of the large variation in values between previous measurements the value is the same as PARP1 association to DNA and set to 1e+5 $M^{-1}s^{-1}$. This rate is dependent on the correction factor $\gamma$ and, thus, the degree of PARP1 PARylation. Corrected for PARP1/DNA concentration interpretation, the default was 1e-2 $M^{-1}s^{-1}$.

**Table S1**. Previously published values of PARPi and PARP1 binding constants used in our models.

| **Binding constants** | | |
|---|---|---|
| PARP1-PARPi kd ($s^{-1}$) | | |
| Veliparib | 4.1E-03 | (*58*) |
| | 7E-03 | (*18*) |
| Niraparib | 5.6E-04 | (*18*) |
| Olaparib | 3.2E-04 | (*18*) |
| Talazoparib | 1.05E-04 | (*58*) |
| | 6.3E-05 | (*18*) |
| PARP1-PARPi ka ($M^{-1}s^{-1}$) | | |
| Veliparib | 1.74E+06 | (*58*) |
| | 1.8E+06 | (*18*) |
| Niraparib | 4.1E+05 | (*18*) |
| Olaparib | 2.3E+05 | (*18*) |
| Talazoparib | 3.68E+05 | (*58*) |
| | 3.6E+05 | (*18*) |

| | | |
|---|---|---|
| PARP1-DNA kd ($s^{-1}$) | | |
| Control | 3.36E-03 | (*17*) |
| | 8.9E-03 | (*18*) |
| Veliparib | 4.08E-03 | (*17*) |
| | 8.5E-03 | (*18*) |
| Niraparib | 4.4E-03 | (*17*) |
| | 7E-03 | (*18*) |
| Rucaparib | 4.77E-03 | (*17*) |
| Olaparib | 2.31E-03 | (*17*) |
| | 1.4E-02 | (*18*) |
| Talazoparib | 2.52E-03 | (*17*) |
| | 5.4E-03 | (*18*) |
| EB-47 | 1.14E-03 | (*17*) |
| UKTT15 | 1.85E-03 | (*17*) |
| BAD | 1.35E-03 | (*17*) |
| Benzamide | 3.28E-03 | (*17*) |
| PARP1-DNA KD (M) | | |
| Control | 3.1E-09 | (*18*) |

| | | |
|---|---|---|
| | ~8E-8 | (*17*) |
| Veliparib | 5.5E-09 | (*18*) |
| | ~3E-7 | (*17*) |
| Niraparib | 8.6E-09 | (*18*) |
| | ~4.8E-7 | (*17*) |
| Rucaparib | ~4.5E-7 | (*17*) |
| Olaparib | 2.4E-09 | (*18*) |
| | ~5E-8 | (*17*) |
| Talazoparib | 4.5E-09 | (*18*) |
| | ~7E-8 | (*17*) |
| PARP1-DNA ka ($M^{-1}s^{-1}$) derived | | |
| Control | 4.2E+4 | (*17*) |
| | 2.9E+6 | (*18*) |
| Veliparib | 1.4E+4 | (*17*) |
| | 1.5E+6 | (*18*) |
| Niraparib | 9.2E+3 | (*17*) |
| | 8.1E+5 | (*18*) |
| Rucaparib | 1.1E+4 | (*17*) |

| | | |
|---|---|---|
| Olaparib | 4.6E+4 | (*17*) |
| | 5.8E+6 | (*18*) |
| Talazoparib | 3.6E+4 | (*17*) |
| | 1.2E+6 | (*18*) |

~ indicates values were interpreted from figures.

**Table S2.** Antibodies used.

| **Antibodies** | | |
|---|---|---|
| PARP1 | Cell Signaling Technology | #9532 |
| PAR | Trevigen | 4336-BPC-100 (discontinued) |
| PAR | Cell Signaling Technology | #83732 |
| γH2aX | Cell Signaling Technology | #2577 |
| Histone H3 | Cell Signaling Technology | #9715 |
| GADPH | R&D Systems | AF5718 |
| RPA1 | Cell Signaling Technology | #2267 |
| BRCA1 | Thermo Fisher | MA1-23164 |
| Dnk pAb to Goat IgG (HRP) | Abcam | Ab97110 |

| Antibodies | | |
|---|---|---|
| Anti-rabbit IgG Fab2 Alexa Fluor 647 | Cell Signaling Technology | #4414 |
| Anti-rabbit IgG, HRP-linked | Cell Signaling Technology | #7074 |
| Anti-mouse IgG, HRP-linked | Thermo Fisher | A-10677 |

**Supplemental Figures**

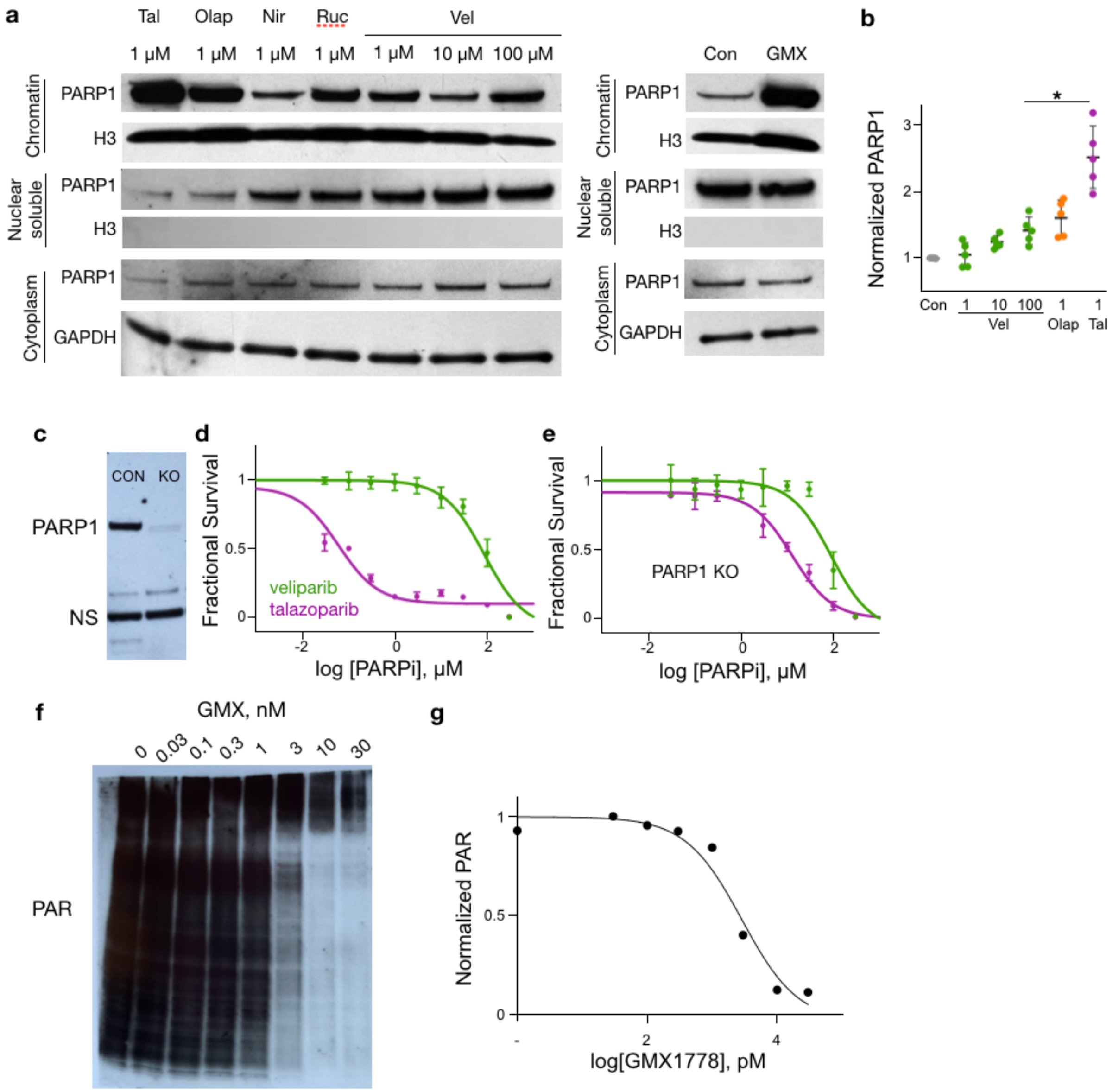


**Fig. S1. PARPi and GMX1778 impact on cells.** (**a**) Representative western blot of HT1080 cells treated with PARPi or GMX1778, H3 – histone H3. Shown are chromatin, nuclear soluble and cytoplasmic fractions. (**b**) Merged results from Fig. 1a and c, n = 5 with average and st. dev., * 100 µM veliparib vs.1 µM talazoparib $p < 0.005$ (Student's t test). (**c**) Western blot of PARP1 expression in normal (con) of PARP1 knockout (KO) cells, non-specific (NS) bands were used as a loading control. (**d**) Dose response of HT1080 cells to treatment with either talazoparib or veliparib. (**e**) Dose response of HT1080 cells with PARP1 knocked out through CRISPR/CAS9 to talazoparib or veliparib. Dose response data average ± st. dev., n=3 biological repeats, with fitted sigmoidal curve. (**f**) PAR western blot and (**g**) quantification of HT1080 cells treated with GMX1778 at varying concentrations for 24 hours followed by treatment with 1 μM $H_2O_2$ for 10 minutes, shown is normalized signal quantification and sigmoidal response curve fit.

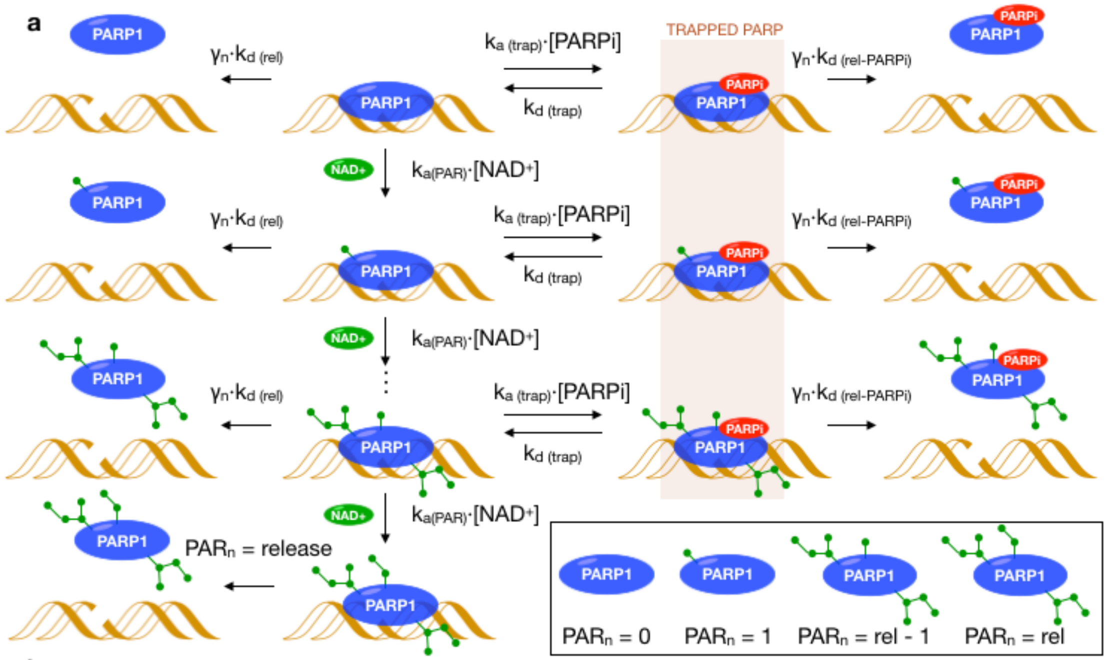


**b**

$$\left(\begin{aligned}
\frac{d(PARP_{DNA})_{PAR_n}}{dt} &= k_{a(PAR)}\cdot[NAD^+]\cdot(PARP_{DNA})_{PAR_{n-1}} - \gamma\cdot k_{d(rel)}\cdot(PARP_{DNA})_{PAR_n} - k_{a(trap)}[PARPi]\cdot(PARP_{DNA})_{PAR_n} + k_{d(trap)}(PARP_{trap})_{PAR_n} - k_{a(PAR)}\cdot[NAD^+]\cdot(PARP_{DNA})_{PAR_n} \\
\frac{d(PARP_{trap})_{PAR_n}}{dt} &= k_{a(trap)}[PARPi]\cdot(PARP_{DNA})_{PAR_n} - k_{d(trap)}(PARP_{trap})_{PAR_n} - \gamma\cdot k_{d(rel\text{-}PARPi)}\cdot(PARP_{trap})_{PAR_n} \\
\frac{d(PARP_{rel\text{-}PARPi})_{PAR_n}}{dt} &= \gamma\cdot k_{d(rel\text{-}PARPi)}\cdot(PARP_{trap})_{PAR_n} \\
\frac{d(PARP_{rel})_{PAR_n}}{dt} &= \gamma\cdot k_{d(rel)}\cdot(PARP_{DNA})_{PAR_n}
\end{aligned}\right)_{n=0\ldots PAR_{release}}$$

**Fig. S2. The ODE model of PARP1-DNA engagement.** (**a**) Model of PARPi induced trapping of PARP1 to damaged DNA. Governed by rate constants, DNA-bound PARP1 either dissociates from DNA, binds $NAD^+$ to undergo ADP-ribosylation, or engages PARPi. PARPi engaged

PARP1 bound to DNA can dissociate from DNA or the PARPi can dissociate. Upon ADP-ribosylation PARP1 is self-PARylated and the model proceeds to $PAR_{n+1}$. At $PAR_n$ = release PARP1 loses all affinity for DNA, by default release = 500. Here, $\gamma_n$ is a correction factor that decreases PARP1 affinity for DNA as a function of PAR level. (**b**) The core differential equations used in the model.

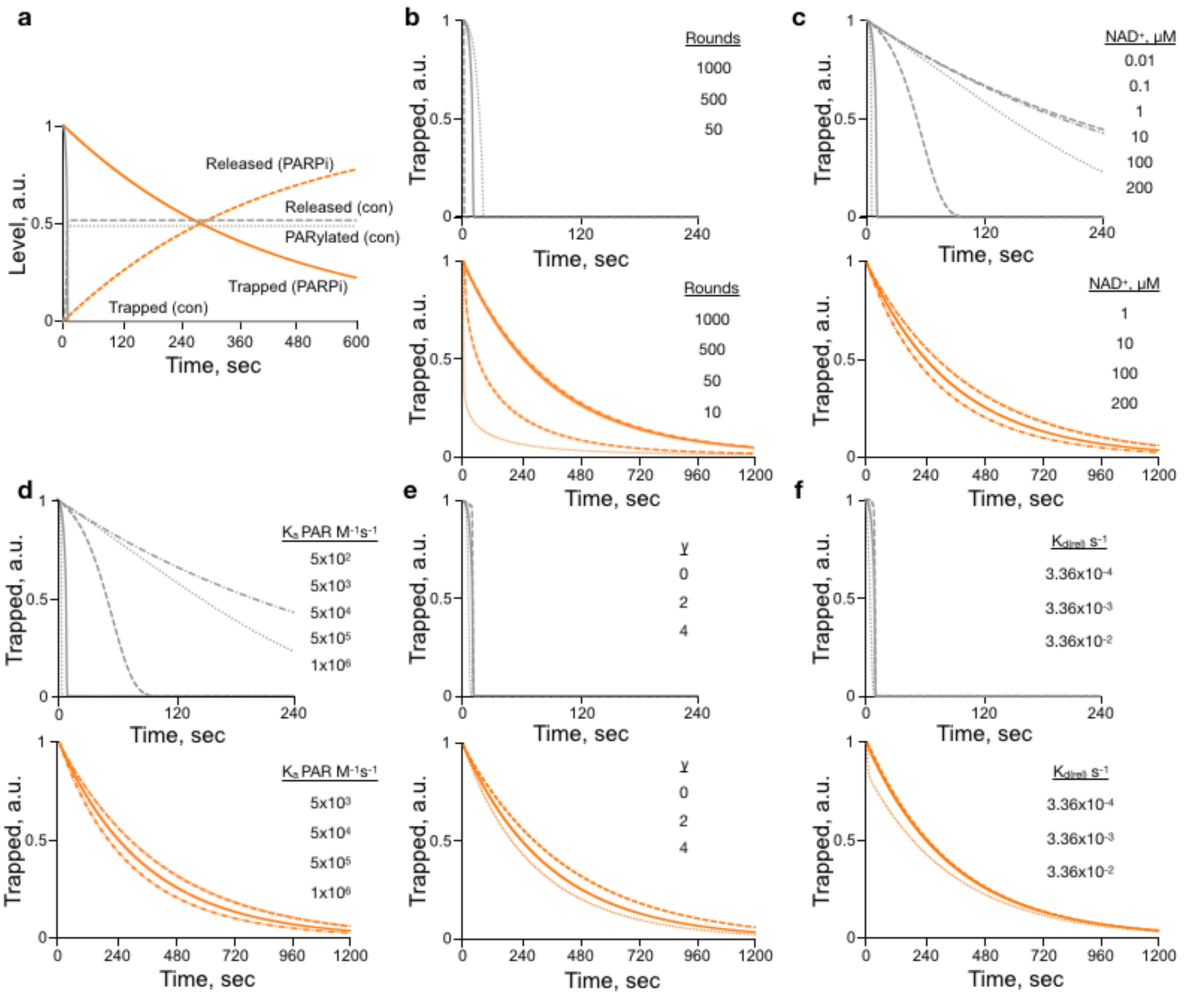


**Fig. S3. The impact of variables on the ODE model of PARP1-DNA interaction.** (**a**) Solution results of species levels in the presence (orange) or absence (gray) of 1 µM olaparib. Trapping (solid line), release (dashed line) and full PARylation (PARylated, dotted line) show the fate of PARP1. In the presence of olaparib all release occurred before full PARylation (500 events) was reached. (**b**) The impact of the number of rounds on the results from the ODE solution shown in the absence (top) and presence (bottom) of 1 μM olaparib. Solid lines are the default value (500). The values in the legend correspond to curves from right to left (high trapping to lower trapping). (**c**) The impact of the $NAD^+$ concentration on the results from the ODE solution shown in the absence (top) and presence (bottom) of 1 μM olaparib. Solid lines are the default value (100 μM). The values in the legend correspond to curves from right to left (high trapping to lower trapping). A lower $NAD^+$ concentration increases trapping by limiting PARylation. (**d**) The impact of the association constant of $NAD^+$ binding to DNA on the results from the ODE solution shown in the absence (top) and presence (bottom) of 1 μM olaparib. Solid lines are the default value ($5x10^5$ $M^{-1}s^{-1}$). The values in the legend correspond to curves from right to left (high trapping to lower trapping). These results share the same impact as $NAD^+$ concentration. (**e**) The impact of $\gamma$ on the results from the ODE solution shown in the absence (top) and presence (bottom) of 1 μM olaparib. Solid lines are the default value (2). The values in the legend correspond to curves from right to left (high trapping to lower trapping). Lower values decrease trapping by lowering the impact of PARylation induced release. (**f**) The impact of the dissociation constant of uninhibited PARP1 from DNA on the results from the ODE solution shown in the absence (top) and presence (bottom) of 1 μM olaparib. Solid lines are the default value ($3.36x10^{-3}$ $s^{-1}$). The values in the legend correspond to curves from right to left (high trapping to lower trapping). Lower values increase trapping for only control by increasing the affinity of PARP1 for DNA.

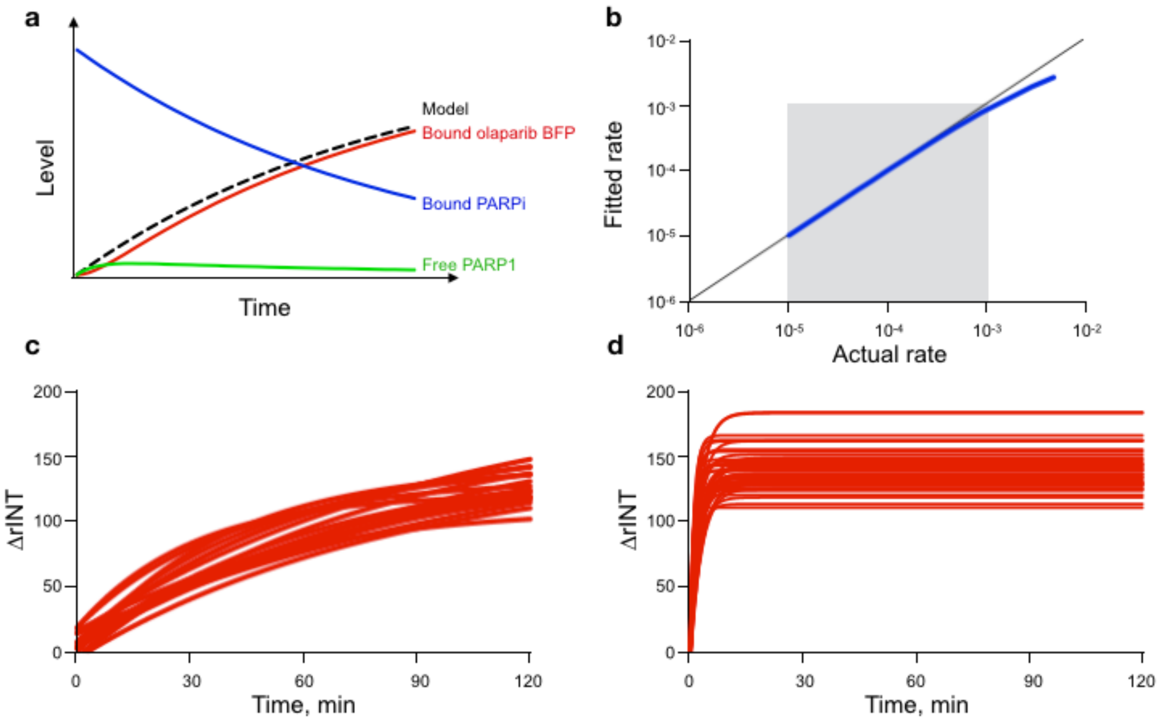


**Fig. S4. Validation of PARPi dissociation measurements.** (**a**) Comparison of fluorescent olaparib binding to PARP1 as PARPi dissociates as approximated by our model (black dashed line) and the ODE solution. Shown are the bound PARPi (blue), free PARP1 (green) and bound fluorescent olaparib (red) as determined by the ODE solution. (**b**) The fitted rate using our model versus actual rate as solved by the ODE equations (blue curve) for the rates under consideration (gray box). At higher dissociation rates the model becomes less accurate as fluorescent drug binding becomes the limiting rate. (**c**) Extended measurements of olaparib dissociation from PARP in HT1080 cells, n = 45 cells, 3 biological repeats. Shown are single cell fitted fluorescent drug association curves demonstrating that shorter experimental times enable accurate rate fitting. (**d**) Binding of fluorescent olaparib in the absence of PARPi in HT1080 cells. Shown are fitted curves for individual cells, n = 32 cell, 3 biological repeats.

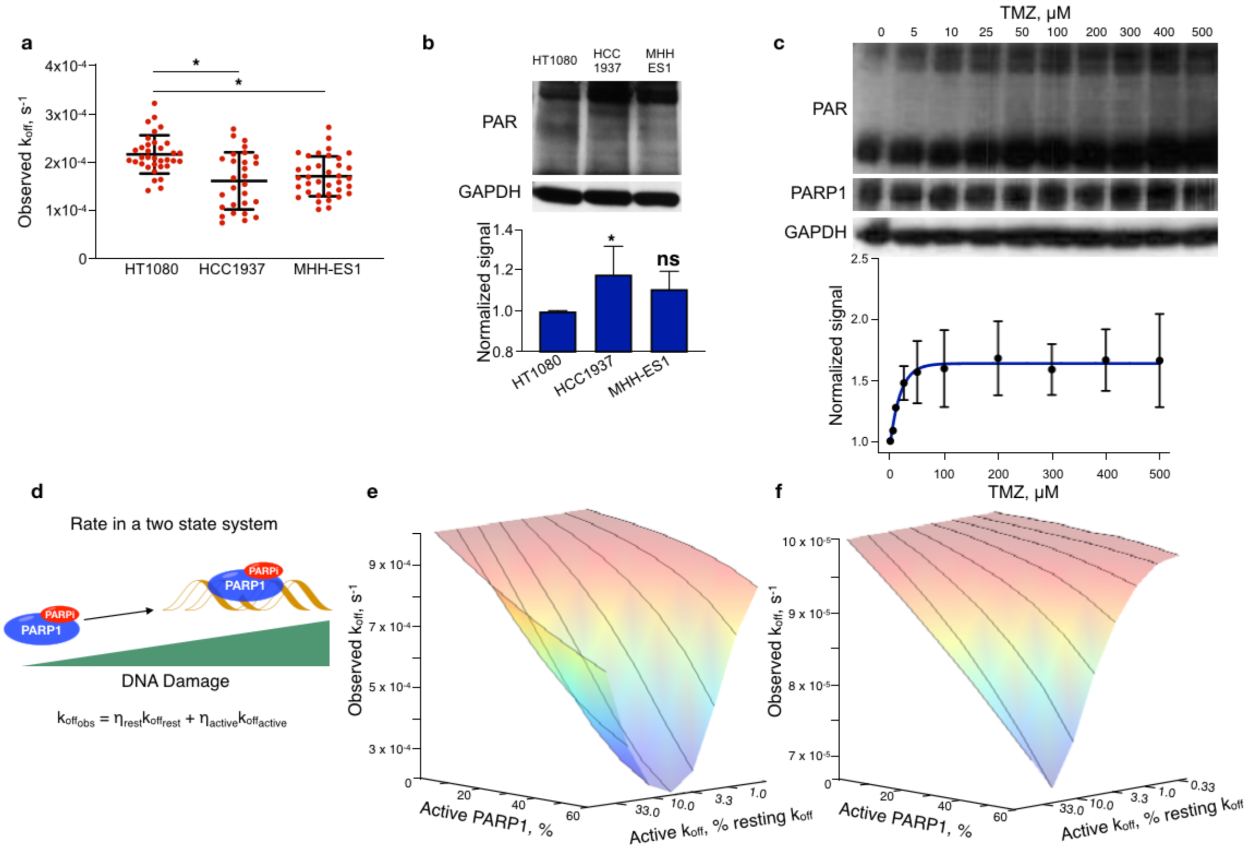


**Fig. S5. Dissociation constant as a function of PAR production.** (**a**) Observed $k_{off}$ of olaparib in three different cell lines. Shown are single cell values with average and st. dev., n >= 28 cells, over 3 experiments for each cell line, *p < 0.001, Student's t test. (**b**). Basal level PAR expression western blot with GAPDH loading controls (top) and quantification (bottom) in HT1080, HCC1937 and MHH-ES1 cell lines. Shown are representative western blot with average and SEM, n=4, *p<0.05 versus HT1080, Student's t test. (**c**) TMZ induced cellular PAR production in HT1080 cells analyzed by western blot, with PARP1 and GAPDH loading controls (top) and quantification (bottom) with one phase association fit curve (blue line). Shown are representative western blot with average and SEM, n=2. (**d**) Depiction of observed $k_{off}$ in a two-state system, where η is the number of molecules in each state. (**e** and **f**) Modeled impact of two $k_{off}$ states on the observed rate as a function of distribution among states and relative $k_{off}$ values.

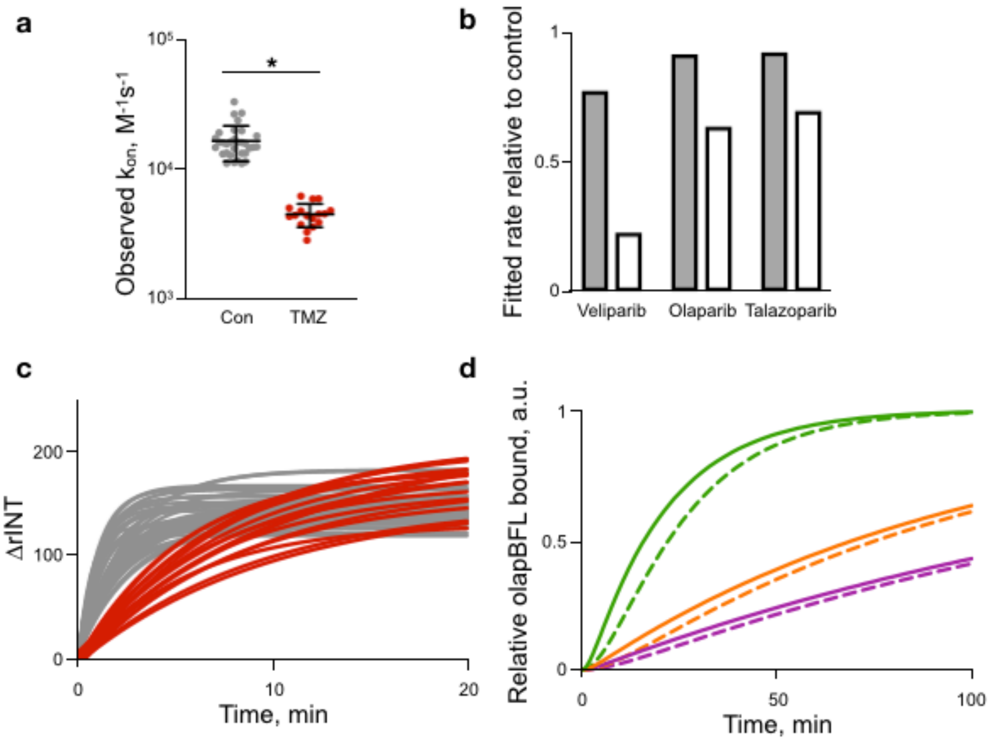


**Fig. S6. Fluorescent olaparib binding in dissociation measurement assay.** (**a**) The observed $k_{on}$ of fluorescent olaparib in in the presence or absence of 100 μM TMZ, shown are single cells with average and st. dev., n >= 17 cells, 3 biological repeats, * $p < 1x10^{-12}$ (Student's t test). (**b**) ODE solved impact of active PARP $k_{on}$ on fitted $k_{off}$ values (gray bars) and the impact of 100 μM TMZ on observed $k_{off}$ (white bars). (**c**) Binding of fluorescent olaparib in the absence of PARPi in untreated HT1080 cells (gray) or cells treated with 100 μM TMZ. Shown are fitted curves for individual cells, n >= 17 cells, 3 biological repeats. (**d**) ODE solution of fluorescent olaparib association to PARP1 in the presence of 1 μM PARPi with $k_{on}$ values measured in control cells (solid line) or cells treated with 100 μM TMZ (dashed lines).

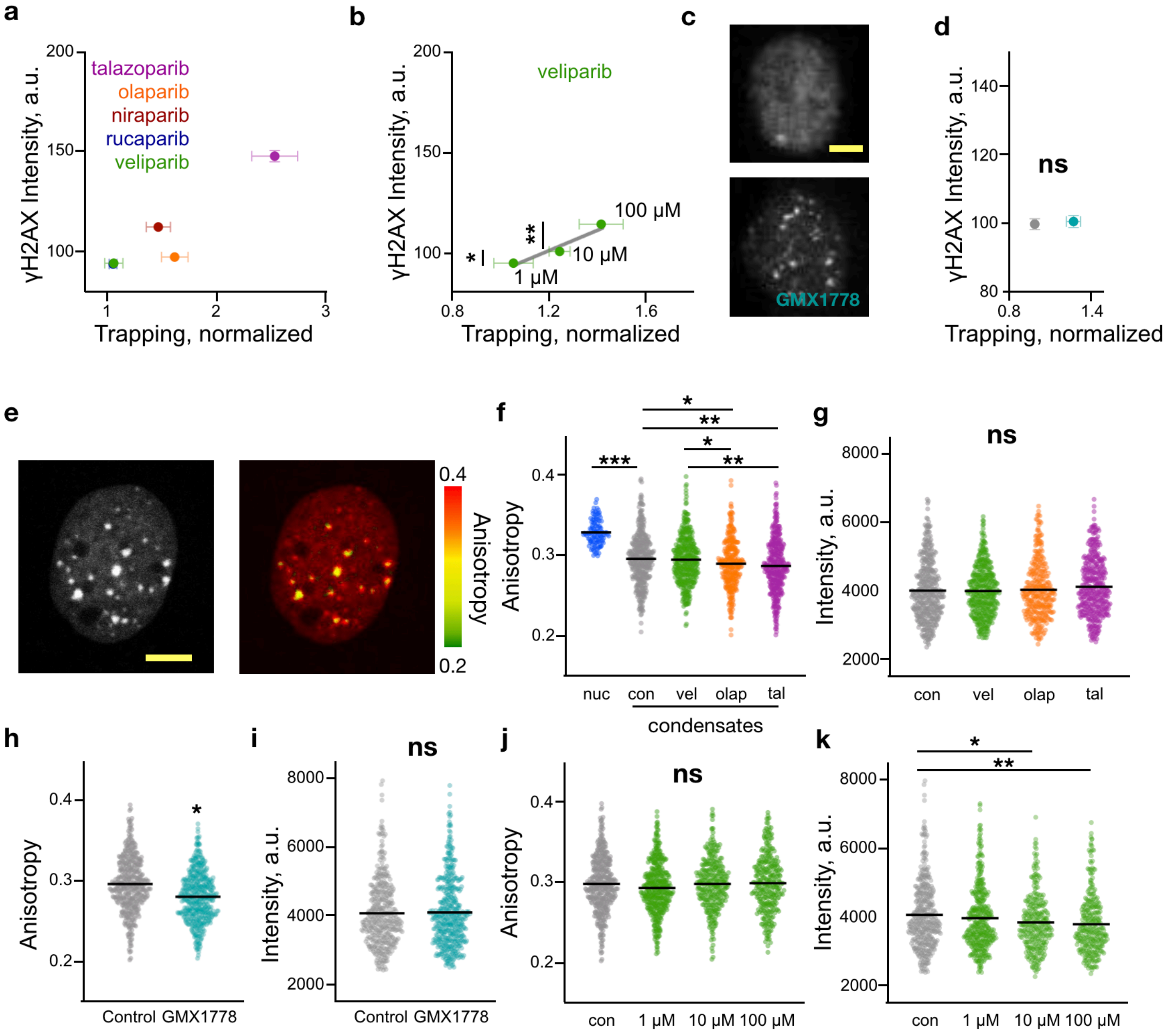


**Fig. S7. PARPi impact on DNA damage induced condensates.** (**a**) Nuclear intensity of γ$H_2$AX IF in HT1080 cells treated with 1 μM PARPi overnight versus PARPi induced PARP trapping, shown are average with SEM, trapping – n=5, DA – n >= 536 cells, 3 biological repeats. (**b**) Nuclear intensity of γ$H_2$AX IF in HT1080 cells treated with veliparib overnight versus PARP1 trapping, with linear fit (gray line), shown are average with SEM, trapping – n=5, DA – n >= 1165 cells, 3 biological repeats, * $p = 3x10^{-3}$, **$p = 2x10^{-8}$ (Student's t test). (**c**) Representative γ$H_2$AX IF images in HT1080 cells untreated or treated with 10 nM GMX1778 for

24 hours (**d**) Nuclear intensity of $\gamma H_2AX$ IF in HT1080 cells treated with 10 nM GMX1778 for 24 hours or control, shown are average with SEM, trapping – n=5, DA – n >= 1285 cells, 3 biological repeats. (**e**) Representative intensity (left) and anisotropy (right) images of an HCC1937 cell nucleus expressing 53BP1-mApple. (**f**) Anisotropy of 53BP1-mApple in nuclei excluding condensates (blue) and condensates in HCC1937 cells treated with 1 μM PARPi or control, and 100 μM TMZ for 1 hour, shown are single cell or condensate values with average (black bar), n = 116 nuclei and n >= 335 condensates, 3 biological repeats, * p < 0.05, ** p < 0.001, *** p < 0.0001 (Student's t test). (**g**) Intensity values of the data in (f). Shown are single condensate values with average (black bar), n >= 335, 3 biological repeats. (**h-i**) Anisotropy (h) and intensity (i) of 53BP1-mApple condensates in HCC1937 cells treated with 10 nM GMX1778 for 24 hours or control, followed by 100 μM TMZ for 1 hour, shown are single condensate values with average (black bar), n >= 405 condensates, 3 biological repeats, * $p < 7x10^{-13}$ (Student's t test). (**j**) Anisotropy and (**k**) intensity values of 53BP1-mApple condensates in HT1080 cells treated with veliparib at difference concentrations or control. Data are single condensates with average, n >= 311 condensates, 3 biological repeats, *p < 0.005, **p=0.0001 (Student's t test). All scale bars = 2 μM.

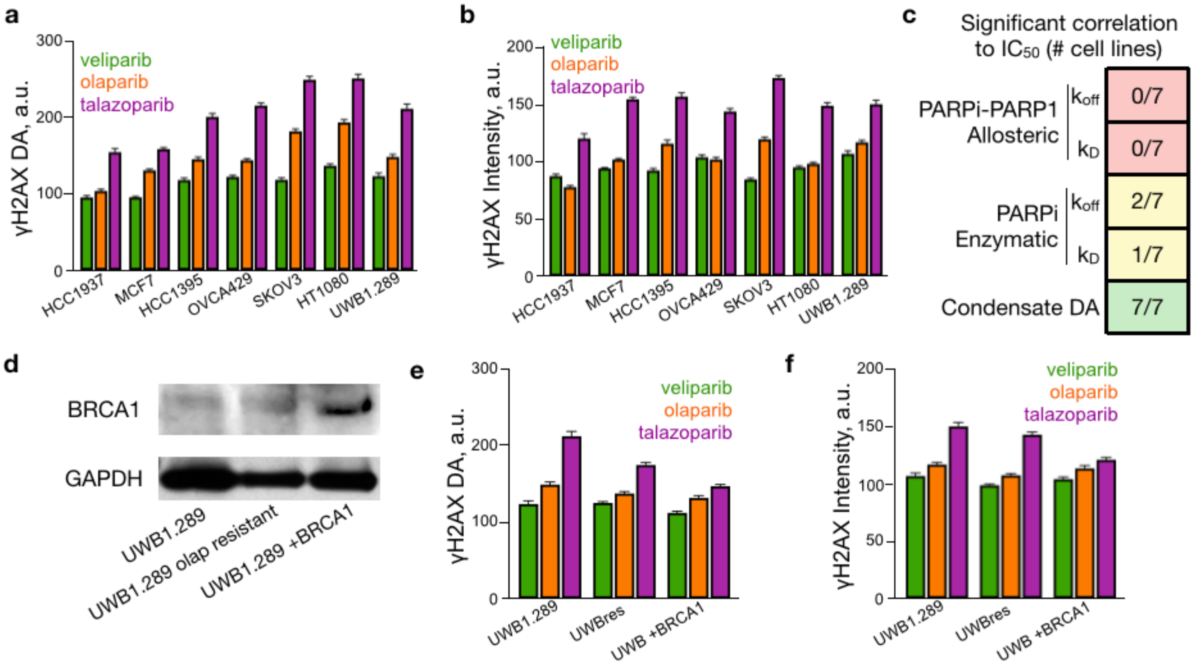


**Fig. S8. PARPi impact on γ$H_2$AX DA in multiple cell lines.** (**a-b**) PARPi specific γ$H_2$AX DA (a) and intensity (b) normalized to control, shown is average with SEM, n >= 536 cells, 3 biological repeats. (**c**) Measured binding property or DA and the number of cell lines with significant correlation, $p < 0.05$ F-test, to the PARPi $IC_{50}$ for talazoparib, olaparib and veliparib. (**d**) BRCA1 expression in UWB1.289, UWB1.289 resistant to olaparib, and UWB1.289 + BRCA1 cells, with GAPDH loading control. (**e-f**) PARPi specific γ$H_2$AX DA (e) and intensity (f) normalized to control, shown is average with SEM, n >= 539 cells, 3 biological repeats.

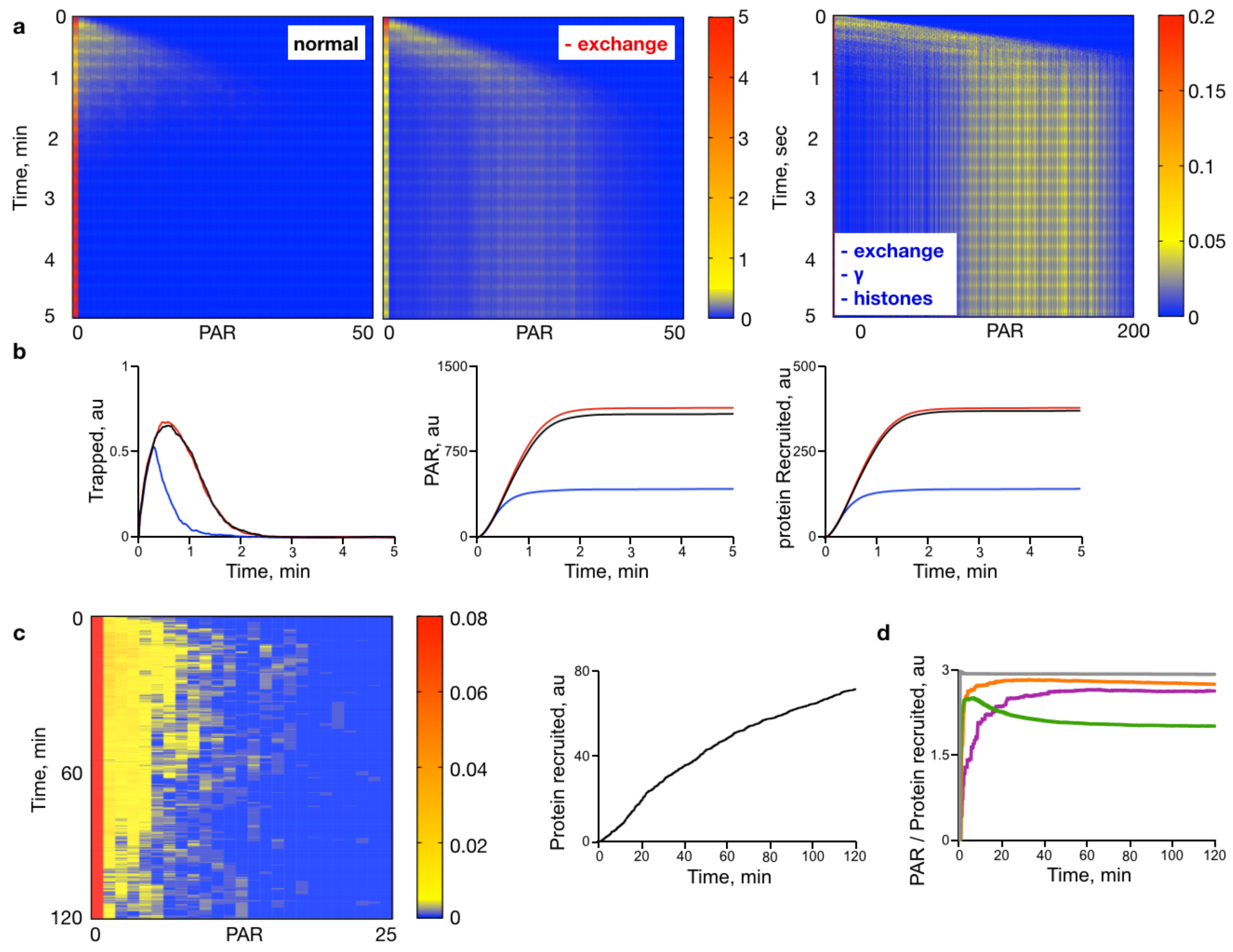


**Fig. S9. Stochastic model results.** (**a**) Heatmap plots of PARP1 PARylation as a function of time. Shown is relative concentration of PARylated PARP1 under normal model conditions (left), in the absence of PARP1 exchange (middle), and in the absence of PARP1 exchange, γ, and histones (right). (**b**) Corresponding trapped PARP1 (left), PAR levels (middle) and protein recruitment (right) to the heatmaps in (a). Shown are simulation results under normal conditions (black), in the absence of PARP1 exchange (red), and in the absence of PARP1 exchange, γ, and histones (blue). (**c**) Heatmap of PARP1 PARylation in the presence of 1 μM olaparib as a function of time (left) and corresponding protein recruitment (right). (**d**) The equilibrium PAR/protein recruited ratio is dependent on the PARPi.

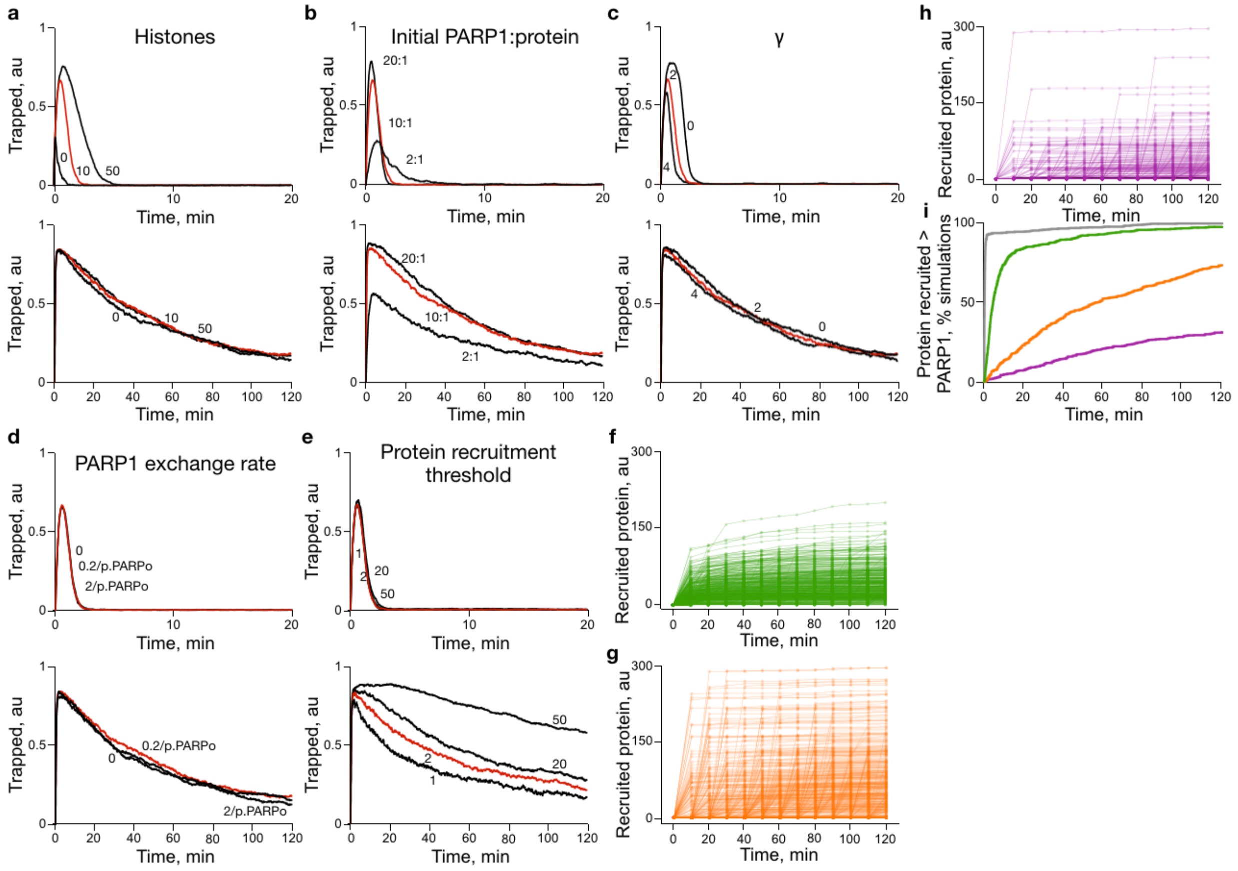


**Fig. S10. Stochastic model tuning and testing.** (**a**) The impact of histone concentration in a biomolecular condensate on the simulation results in the absence (top) and presence (bottom) of 1 μM olaparib. Red lines are the default value (5). (**b**) The impact of the ratio of PARP1 to initial other DNA damage binding proteins in a biomolecular condensate on the simulation results in the absence (top) and presence (bottom) of 1 μM olaparib. Red lines are the default value (10:1). (**c**) The impact of the PARP1 DNA affinity PARylation dependence factor γ on the simulation results in the absence (top) and presence (bottom) of 1 μM olaparib. Red lines are the default value (2). (**d**) The impact of the exchange rate of PARylated PARP1 in a biomolecular condensate exchange with unPARylated PARP1 on the simulation results in the absence (top) and presence (bottom) of 1 μM olaparib. Red lines are the default value (0.2/p.PARP0). (**e**) The impact of the DNA binding protein recruitment threshold value on the simulation results in the absence (top) and presence (bottom) of 1 μM olaparib. Red lines are the default value (2). (**f-h**) Single simulation values of recruited protein as a function of time in the presences of 1 μM (f) veliparib, (g) olaparib, and (h) talazoparib. (**i**) Stochastic simulation results of protein recruited to levels greater than PARP1 in the presence or absence of 1 μM PARPi. All tracings (except f-h) are an average of 1000 simulations.

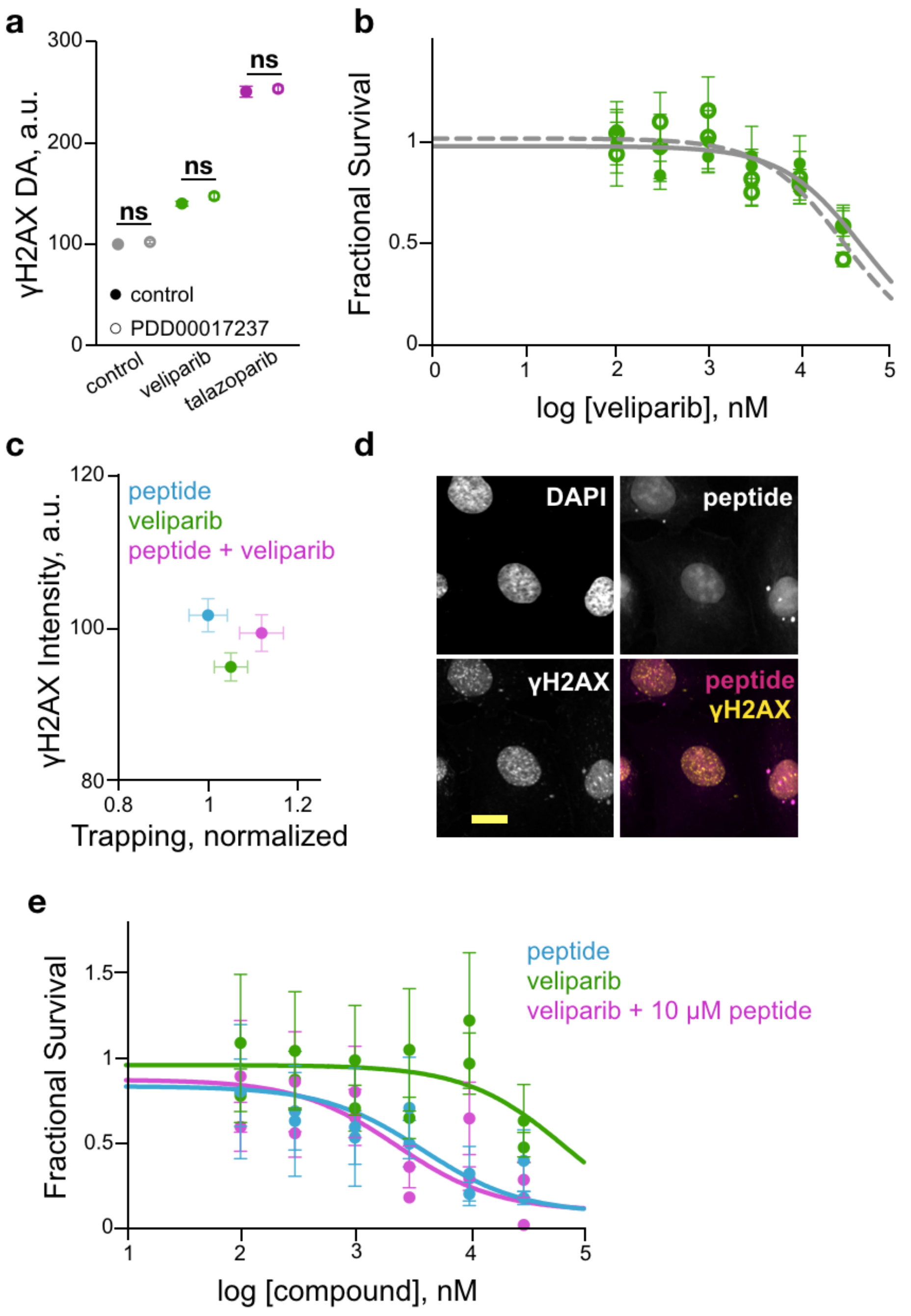


**Fig. S11. Impact of altered PAR availability on trapping.** (**a**) γ$H_2$AX DA in HT1080 cells treated in the absence of PARPi (control), or with 1 µM veliparib or 1 µM talazoparib with or without 100 nM PDD00017273 overnight, shown are average with SEM, n >= 378 cells. There were no significant differences between cells with and without PDD00017273, Student's t test. (**b**) Dose response of HT1080 cells to veliparib in the absence (closed circles – solid line) or presence (open circles – dashed line) of 100 nM PDD00017273, shown are average with standard deviation of two experiments, n = 3 for each experiment, and sigmoidal curve fit (prism) to the average of the two experiments. (**c**) Nuclear intensity of γ$H_2$AX IF in HT1080 cells treated with 1 μM veliparib, 10 μM PAR binding peptide or the combination overnight versus PARPi induced PARP trapping, shown are average with SEM normalized to control, trapping - n=5, DA - n >= 917 cells, 3 biological repeats, no condition showed a significant increase over control (Student's t test). (**d**) Representative images of HT1080 cells treated overnight with 10 μM of PAR binding peptide, fixed and immunolabeled for γ$H_2$AX and stained with DAPI, scale bar = 5 μm.

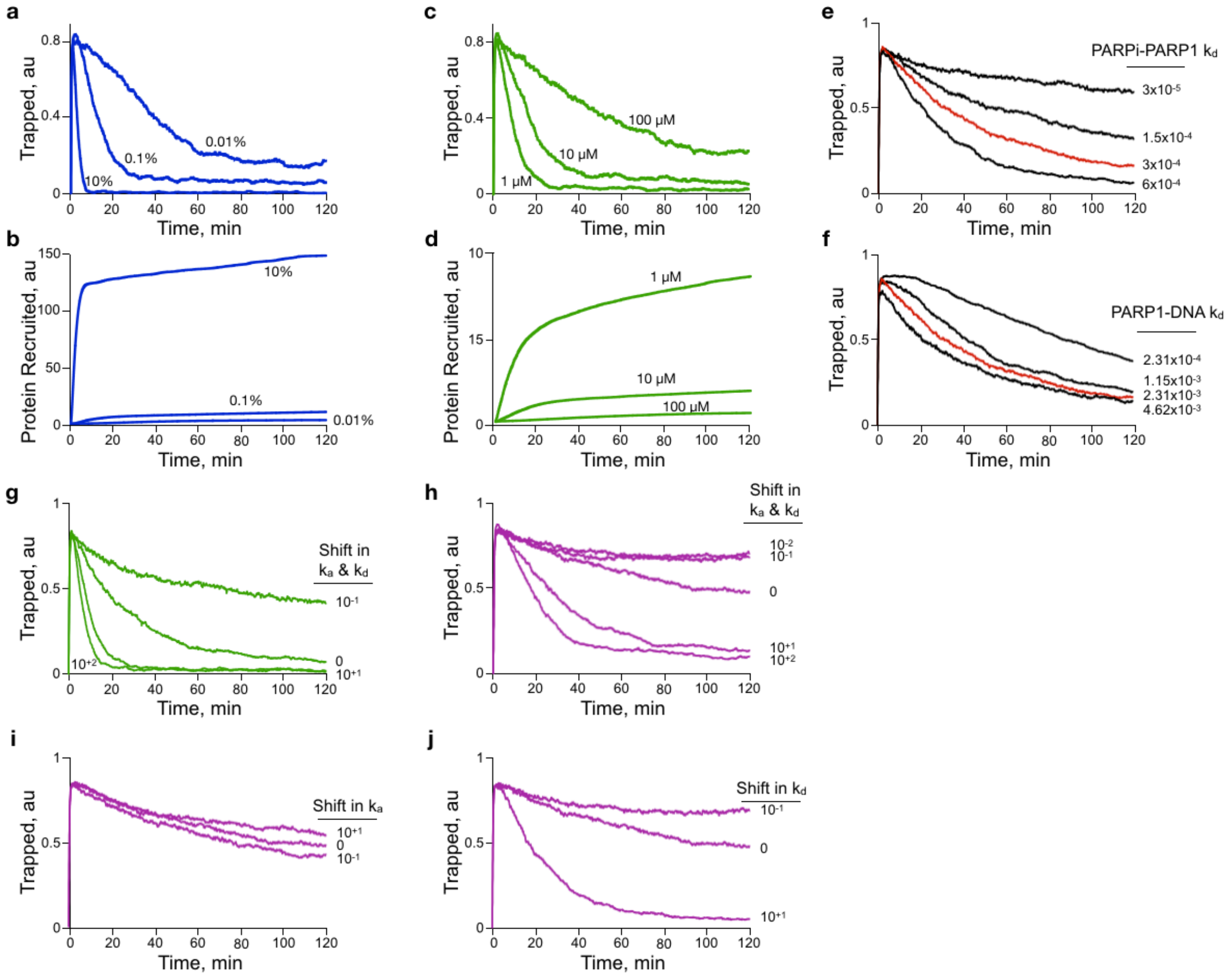


**Fig. S12. Modeling the impact of different rate constants.** (**a-b**) Stochastic simulation results of trapped PARP1 on DNA (a) and condensate recruited DDR protein levels (b) as a function of time at decreasing nuclear $NAD^+$ concentration levels, shown are the percentage of baseline $NAD^+$ concentration. (**c-d**) Stochastic simulation results of trapped PARP1 on DNA (c) and condensate recruited DDR protein levels (d) as a function of time at 3 different veliparib concentrations. (**e**) The impact of synthetic PARPi dissociation constant (*p.kd_trap*) values on trapping within a biomolecular condensate. Shown are simulation results with the labeled $k_d$ value and default olaparib values for other binding constants and 1 μM olaparib. The measured value used in normal conditions is $3x10^{-4}$ $s^{-1}$ (red line). (**f**) The impact of synthetic olaparib bound PARP1 dissociation from DNA constant (*p.kd_relD0*) values on trapping within a biomolecular condensate. Shown are simulation results with the labeled $k_d$ values and default olaparib values for other binding constants and 1 μM olaparib. The measured value used in normal conditions is $2.31x10^{-3}$ $s^{-1}$ (red line). (**g**) The impact of synthetic veliparib at 1 μM. Shown are normal (0 shift) and shifts in both $k_a$ and $k_d$ (the equilibrium binding constant $k_D$ remains the same). (**h**) The impact of synthetic talazoparib at 1 μM. Shown are normal (0 shift) and shifts in both $k_a$ and $k_d$ (the equilibrium binding constant $k_D$ remains the same). (**i**) The impact of synthetic talazoparib at 1 μM. Shown are normal (0 shift) and shifts in only $k_a$, which alters the equilibrium binding constant $k_D$ by the same degree. (**j**) The impact of synthetic talazoparib at 1 μM. Shown are normal (0 shift) and shifts in only $k_d$, which alters the equilibrium binding constant $k_D$ by the same degree. All tracings are an average of 1000 simulations.

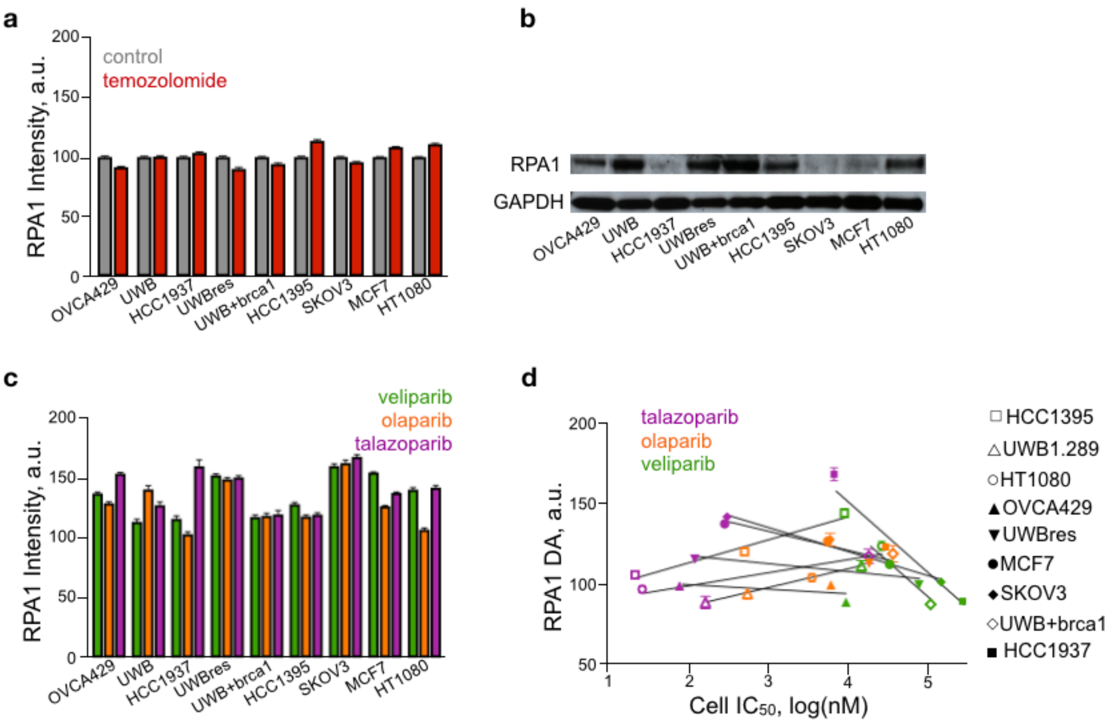

**Fig. S13. RPA1 condensate recruitment.** (**a**) RPA1 nuclear intensity in cells untreated or treated with 100 µM TMZ overnight, shown are average with SEM, n >= 680 cells, 3 biological repeats, normalized to control for each cell line. (**b**) Western blot RPA1 expression levels for each cell line in (a) along with GAPDH loading controls. (**c**) RPA1 nuclear intensity in cells treated with 1 µM PARPi overnight, shown are average, normalized to untreated control, with SEM, n >= 350 cells, 2 biological repeats. (**d**) RPA1 DA as a function of cell line IC50 for each cell line and PARPi shown in Fig. 5b. Shown are average, normalized to untreated control, with SEM, n >= 350 cells, 2 biological repeats and linear fit (prism) for each cell line.
